# Emergence and rapid dissemination of highly pathogenic avian influenza virus H5N1 clade 2.3.4.4b in wild birds, Chile

**DOI:** 10.1101/2023.04.07.535949

**Authors:** Naomi Ariyama, Catalina Pardo-Roa, Gabriela Muñoz, Carolina Aguayo, Claudia Ávila, Christian Mathieu, Barbara Brito, Rafael Medina, Magdalena Johow, Victor Neira

**Affiliations:** Departamento de Medicina Preventiva Animal, Facultad de Ciencias Veterinarias y Pecuarias, Universidad de Chile. 11735 Santa Rosa, La Pintana, Santiago, Chile. (N. Ariyama, G. Muñoz, V. Neira); Departamento de Enfermedades Infecciosas e Inmunología Pediátrica, Escuela de Medicina, Pontificia Universidad Católica de Chile, Santiago, Chile. (C. Pardo-Roa, R. Medina); Servicio Agrícola y Ganadero (SAG), Santiago, Chile. (C. Aguayo, C. Ávila, C. Mathieu, M. Johow); School of Nursing, Pontificia Universidad Católica de Chile, Santiago, Chile. (C. Pardo-Roa); Veterinary Research Office, Department of Primary Industries, Menangle, New South Wales, Australia (B. Brito).; Department of Pathology and Laboratory Medicine, School of Medicine, Emory Vaccine Center, Emory University, Atlanta, USA (R. Medina); Department of Microbiology, Icahn School of Medicine at Mount Sinai, New York, USA (R. Medina).

**Keywords:** Avian Influenza virus, Influenza A Virus, H5N1 Subtype, Wild birds, Viral dissemination.

## Abstract

In December 2022, HPAI H5N1 clade 2.3.4.4b emerged in Chile. We detected the virus in 93 wild bird samples and sequenced the whole genome of nine Chilean strains from pelicans and gulls. Phylogenetic analysis suggests at least two different HPAI viral clusters in South America.

---

Highly pathogenic avian influenza (HPAI) viruses H5N1 subtype grouped within the HA clade 2.3.4.4b, are spreading globally and causing high mortalities in domestic and wild birds (1). These viruses had spilled over to several non-avian species, including humans (2). To contain the outbreaks culling HPAI-infected or exposed poultry has resulted in the disposal of around 120 million domestic birds in 2022 (3). Therefore, HPAI poses a threat not only to public health due to its zoonotic potential but also to food security.

In late 2021, the H5N1 clade 2.3.4.4b, which spread predominantly in Europe, Asia, and Africa, was detected in wild birds in North America and shortly after in poultry (3–5). In October 2022, HPAI reached South America and was officially reported for the first time in Colombia and later in Peru, Ecuador, and Venezuela (2).

In early December 2022, increased deaths among wild birds were detected across the north coast of Chile (Appendix Figure 1). Wild birds, mainly pelicans, were found dead or dying. Until December 22, 2022, the official Veterinary Services of the Chilean Agricultural and Livestock Service (SAG) had collected 1368 samples from domestic (n=1080) and wild birds (n=288) for HPAI detection and epidemiological investigation (Appendix Table 1). A total of 578 real-time RT-PCR (RT-qPCR) reactions and 790 agar gel immunodiffusion tests (AGID) were performed to detect active infection and previous exposure to the virus.

The RT-qPCR was performed with the VetMAX-Gold AIV Detection Kit (Applied Biosystems™), targeting the avian influenza virus (AIV) M gene. Samples included: tissues (n= 13, 15% of positivity), tracheal (n= 248, 17%), cloacal (n= 314, 15%), and oral swabs (n= 3, 33%). A total of 93 samples from Arica y Parinacota (n=18), Tarapaca (n=18), Antofagasta (n=53), and Atacama (n=4) regions were considered AIV-positive qRT-PCR (Ct value <40, 16 % of positivity) (Appendix Figure 1). No poultry samples were positive to AIV. The most frequently AIV-infected species were pelicans (*Pelecanus thagus*, n=50, 54%), followed by vultures (*Cathartes aura*) and Peruvian boobies (*Sula variegata)* (Appendix Table 1).

All serum samples collected mainly from domestic birds (n= 785 out of 790) and processed using the official AGID protocol tested negative for AIV (6). Next, full-genome sequencing was attempted in 11 RT-qPCR-positive samples selected by Ct value and location. The eight segments of AIV genome were amplified using the MS RT-PCR approach (7). Then they were sequenced using the whole-genome amplicon-based sequencing protocol by Oxford Nanopore Technology according to the manufacturer’s instruction (Native Barcoding Kit 96, SQK-NBD114.96). Genome assembly was performed using the genome A/Falco_rusticolus/EdoMex/CPA-19638-22/2022(H5N1) (GenBank accession numbers OP691321 to OP691328) as a reference due to its highest similarity our *de novo* assemblies. Ten samples were sequenced with a mean coverage depth of 33.381x, and 9 of 10 complete genome were obtained (Table) (Accession numbers OQ455414 - OQ455462).

All samples were classified as H5N1 subtype and 2.3.4.4b H5 clade (Appendix). A/Peru/LIM-003/2022 and A/Peru/LAM-002/2022 (GISAID Isolate IDs EPI_ISL_16249730 and EPI_ISL_16249681) were the most closely related HA sequences found on the GISAID database (8) (Figure A). Similar results were observed on the NA (Figure B) and internal genes phylogenies (Appendix Figures 2-7). The Peruvian sequences correspond to isolates collected in November 2022 from domestic chickens. Sequences from Ecuador and Mexico were observed in the same subcluster when enough-quality sequences (n content < 0.5%) were available for the analyses (Appendix Figures 2-7). For HA, the Chile-Peru subcluster showed a non-synonymous mutation T392A (L131Q), previously associated with antigenic variability in H5N1 strains (9). Other synonymous and non-synonymous mutations, which are not associated with phenotypic changes to date, were found in NA (L269M and S339P) and the internal genes (Appendix Table 2). Sequences from Venezuela grouped outside the Chile-Peru subcluster for all its available complete sequences (H5, N1, M, and PA), suggesting a different introduction to South America. However, due to the low availability of HPAI H5N1 sequences from Central and South America, conclusions on the origin of the Chilean clusters are limited. Our analysis supports the idea of the HPAI H5N1 clade 2.3.4.4b introduction to the Americas across the Atlantic flyway and its posterior dissemination to all the other flyways (https://www.usgs.gov/centers/nwhc/science/distribution-highly-pathogenic-avian-influenza-north-america-20212022). These flyways reach the southernmost tip of the South American mainland, representing a high risk as an HPAI dissemination route across the continent (10). Thus, suggesting the prompt arrival of the HPAIV to the Austral region and hazardous proximity to Antarctica.

According to official data, until January 18, 2023, the HPAI virus H5N1 has disseminated as far as the Maule region, and no poultry cases have been confirmed. The SAG has implemented a contingency plan to perform extensive surveillance and reinforcement of biosecurity measurements to avoid the introduction of the HPAIV into poultry. Further studies are needed to determine the impact of H5N1 HPAIV in the country and the potential introduction to Antarctica.

## Acknowledgments

We thank the Agricultural and Livestock Service (SAG) personnel for their support and contributions, especially in the sample collection. We are grateful to Belen Aguero and Felipe Berrios from the Animal Virology Lab, Universidad de Chile, for their help in sample processing. Thank to Leonardo Almonacid at Unidad de Bioinformática y Biología Computacional Pontificia Universidad Católica de Chile for his support on the bioinformatics pipeline. We are grateful to the GISAID EpiFlu™ Database, laboratories, and source of original data of influenza A virus (IAV) sequences, especially to the Servicio Nacional de Sanidad Agraria del Perú – SENASA, Peru, source of the strain closest strains.

## Author Bio

Naomi Ariyama is DVM and Ph.D. student at the University of Chile; her primary focus is the study of emerging viral zoonotic pathogens. Catalina Pardo-Roa DVM, MSc, and Ph.D. is an associate investigator at the Molecular Virology Laboratory; her focus is the study of IAV and NGS.

---

**Table.** Summary of sequencing results of highly pathogenic avian influenza virus H5N1 clade 2.3.4.4b in wild birds, December 2022, Chile.

| Strain | Species | Region | Sample type | Collection Date | Ct* | Total reads | Influenza like reads (%) | Coverage (average) |
| --- | --- | --- | --- | --- | --- | --- | --- | --- |
| 226924-1 | Pelican | Arica y Parinacota | Tracheal swab | 12/6/2022 | 31 | 839824 | 716890 (92) | 47593x |
| 226618-1 | Pelican | Arica y Parinacota | Tracheal swab | 12/5/2022 | 26 | 170767 | 144419 (92) | 11717x |
| 226618-2 | Pelican | Arica y Parinacota | Cloacal swab | 12/5/2022 | 26 | 305220 | 255977 (92) | 20429x |
| 226955-1 | Pelican | Antofagasta | Tracheal swab | 12/6/2022 | 19 | 239414 | 193780 (87) | 8700x |
| 226955-3 | Pelican | Antofagasta | Tracheal swab | 12/6/2022 | 19 | 215419 | 166728 (84) | 8300x |
| 226958-1 | Pelican | Antofagasta | Tracheal swab | 12/6/2022 | 17 | 552755 | 387062 (77) | 22680x |
| 227023-1 | Pelican | Tarapaca | Cloacal swab | 12/7/2022 | 25 | 868 | 245 (81) | 24x |
| 227023-2 | Belcher's gull | Tarapaca | Cloacal swab | 12/7/2022 | 24 | 678724 | 594285 (95) | 63145x |
| 227023-3 | Belcher's gull | Tarapaca | Tracheal swab | 12/7/2022 | 21 | 1809324 | 1593584 (95) | 151985x |
| 227087-1 | Pelican | Arica y Parinacota | Tracheal swab | 12/7/2022 | 25 | 19121 | 15983 (92) | 1237x |

\* Cycle threshold of the real time RT-PCR for the matrix gene of influenza A virus.

**Figure.**
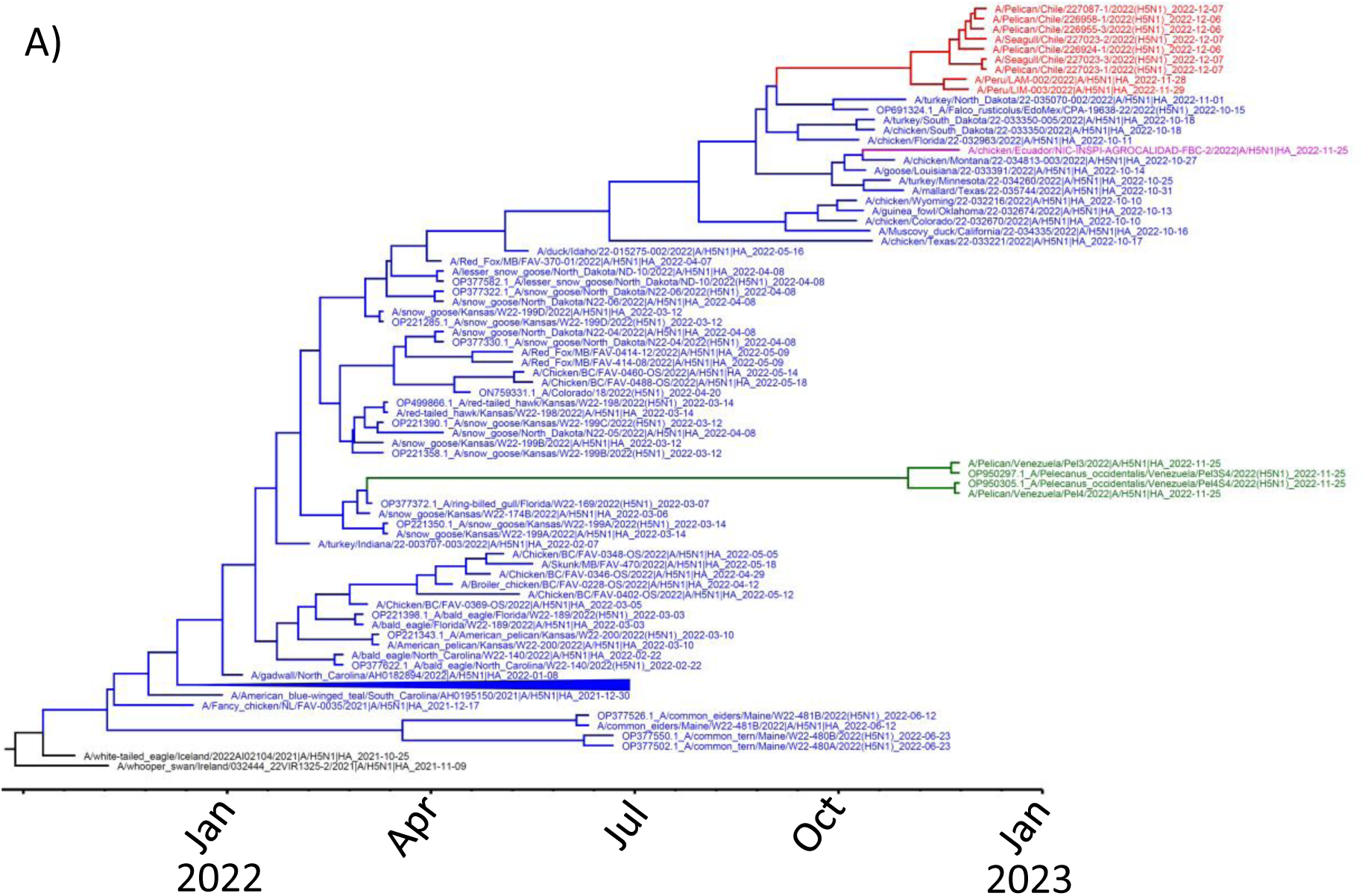
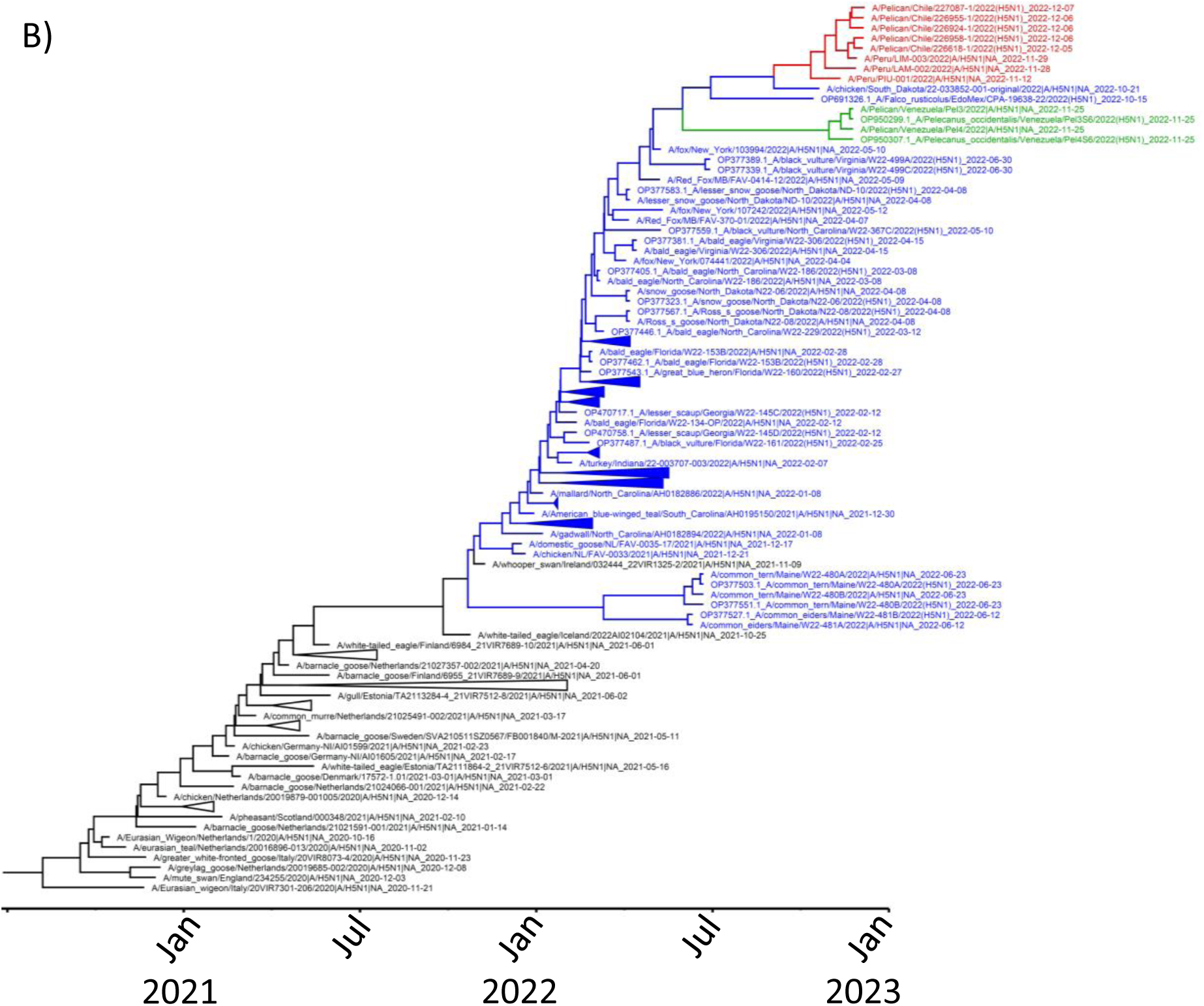
A) Time-scale Bayesian MCC tree of 2.3.4.4b H5 clade. B) Time-scale Bayesian MCC tree of N1 from isolates of H5N1 subtype 2.3.4.4b H5 clade. The Chilean-Peruvian subcluster is highlighted in red, the Venezuelan strains in green, Ecuadorian in pink, the North American in blue, and other reference sequences in black.

**Appendix Table 1.**
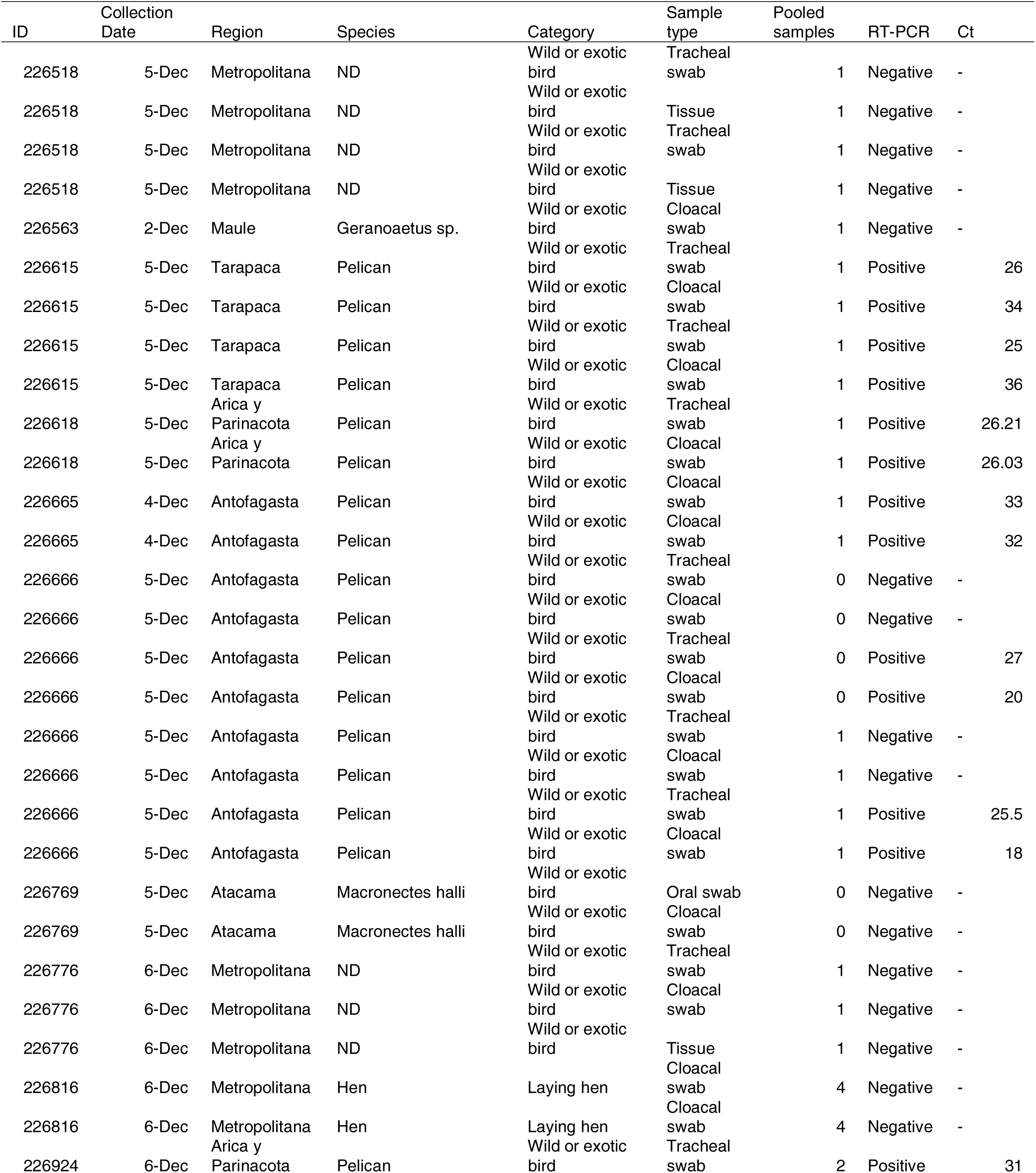
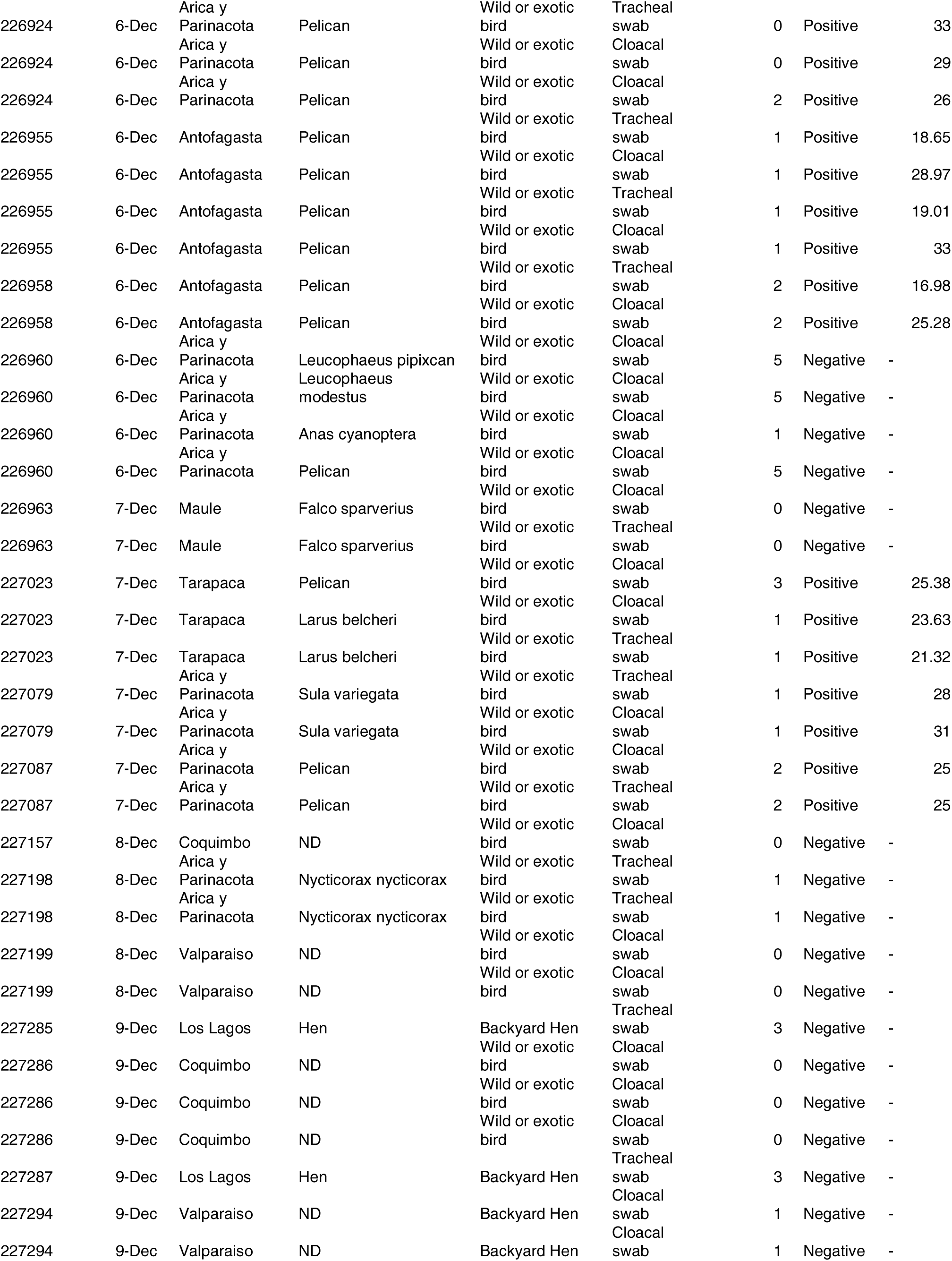
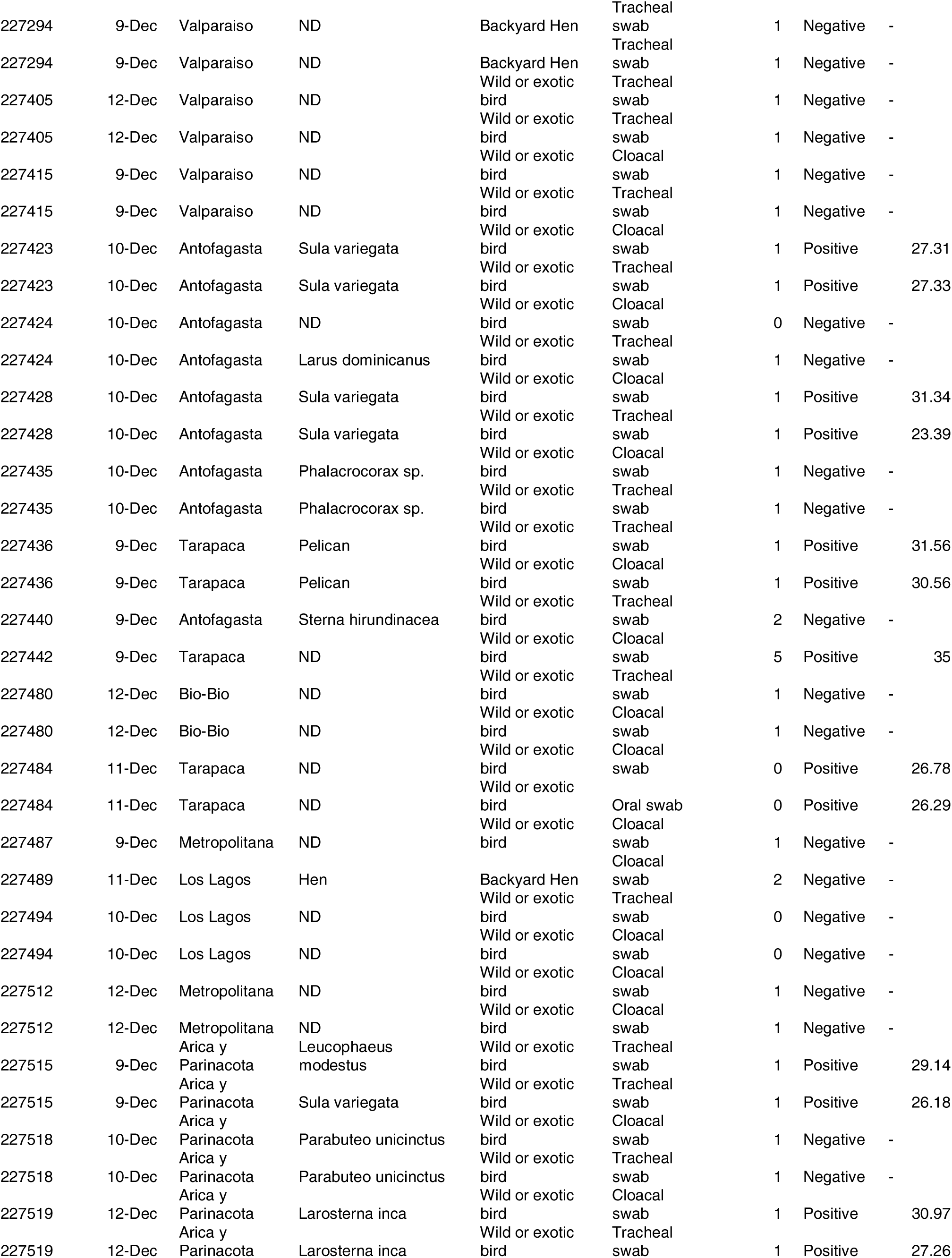
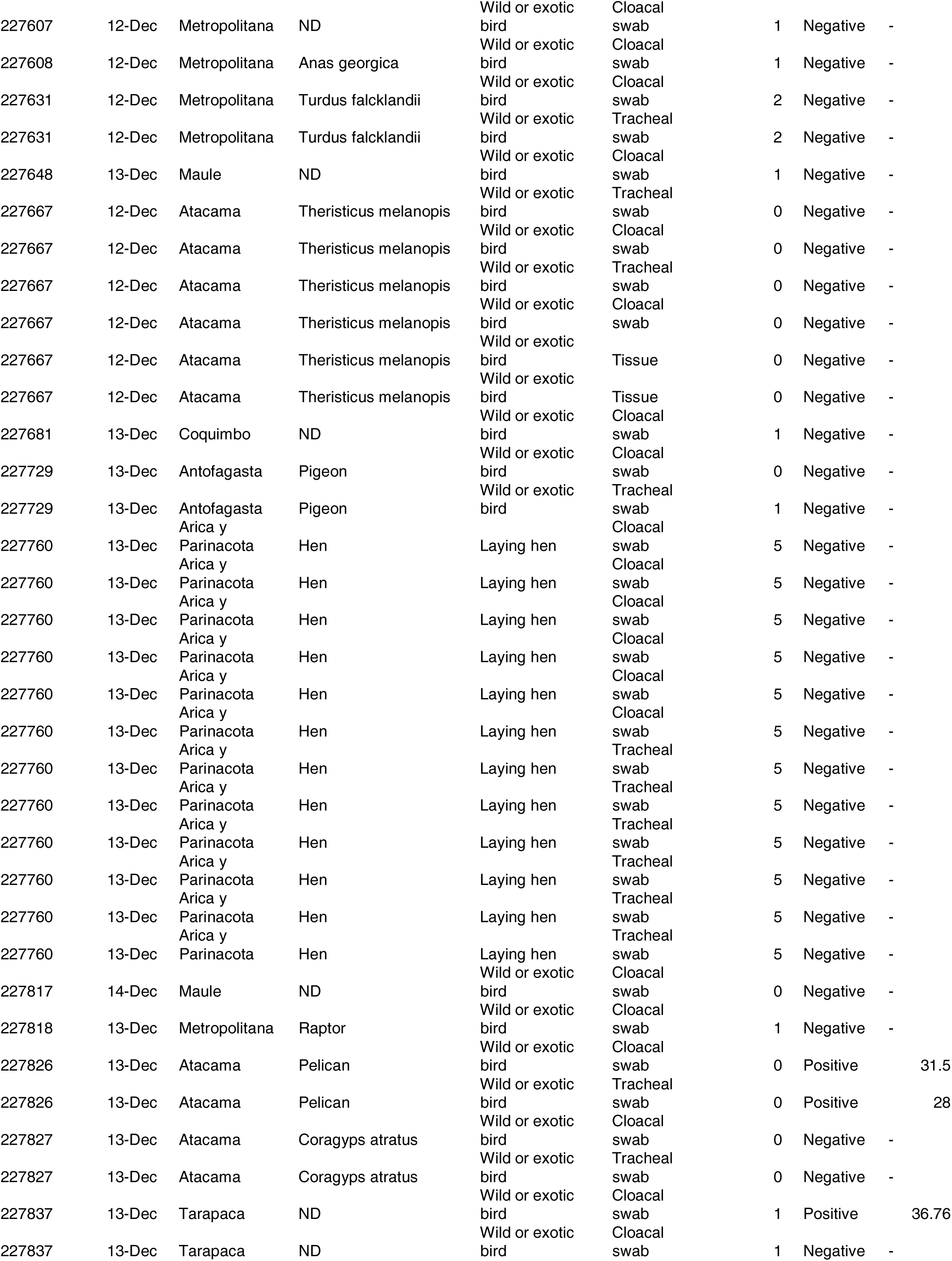
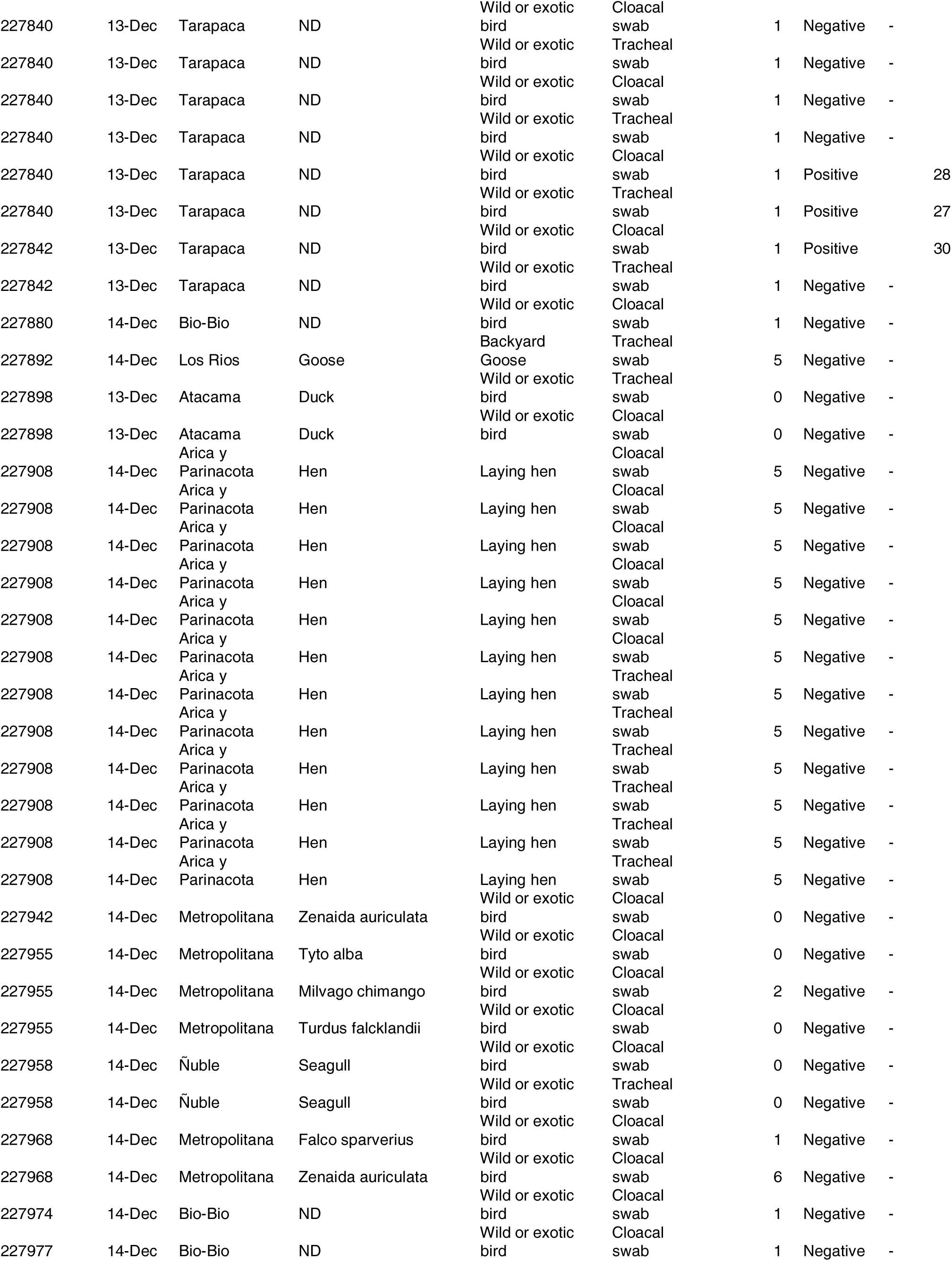
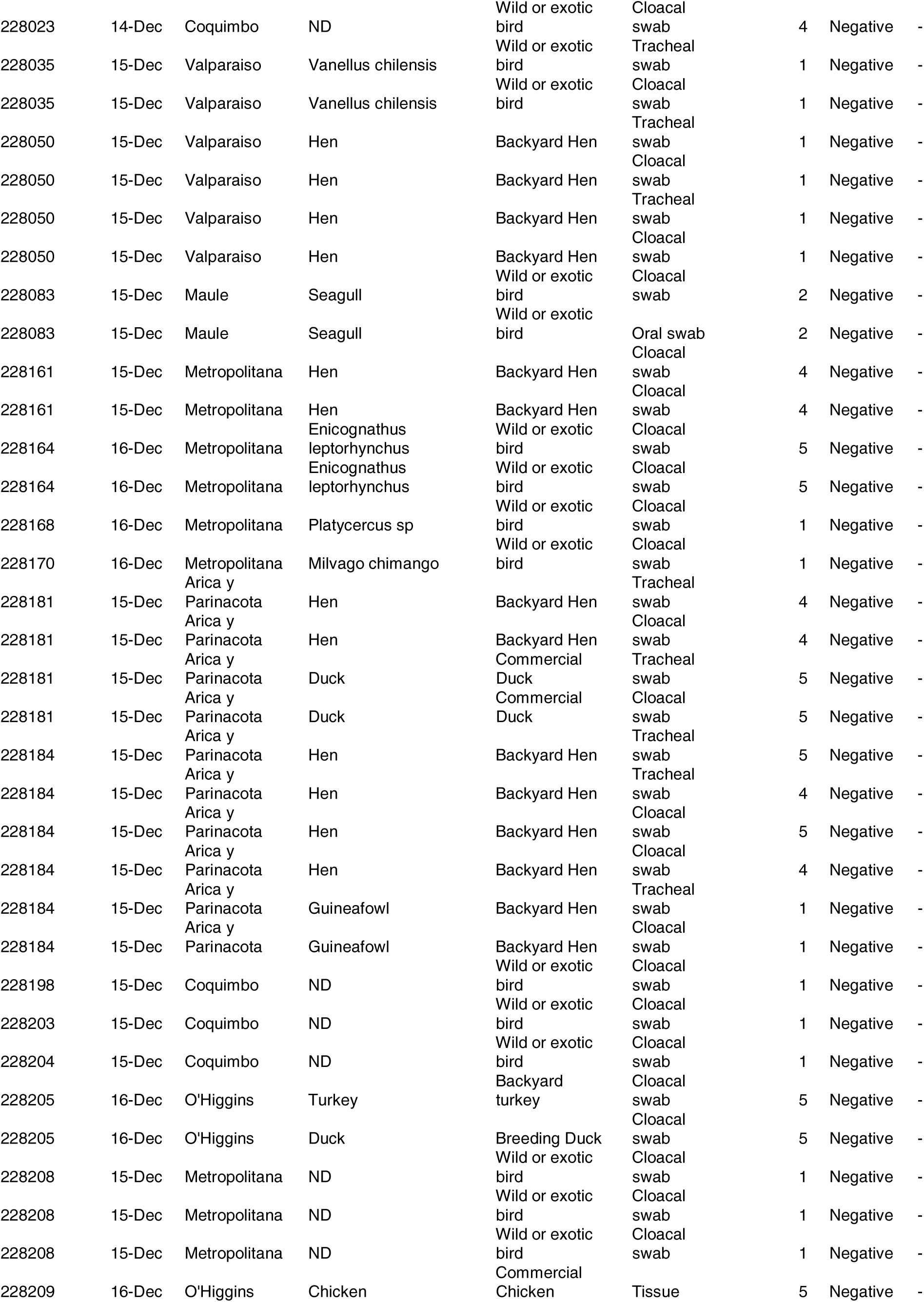
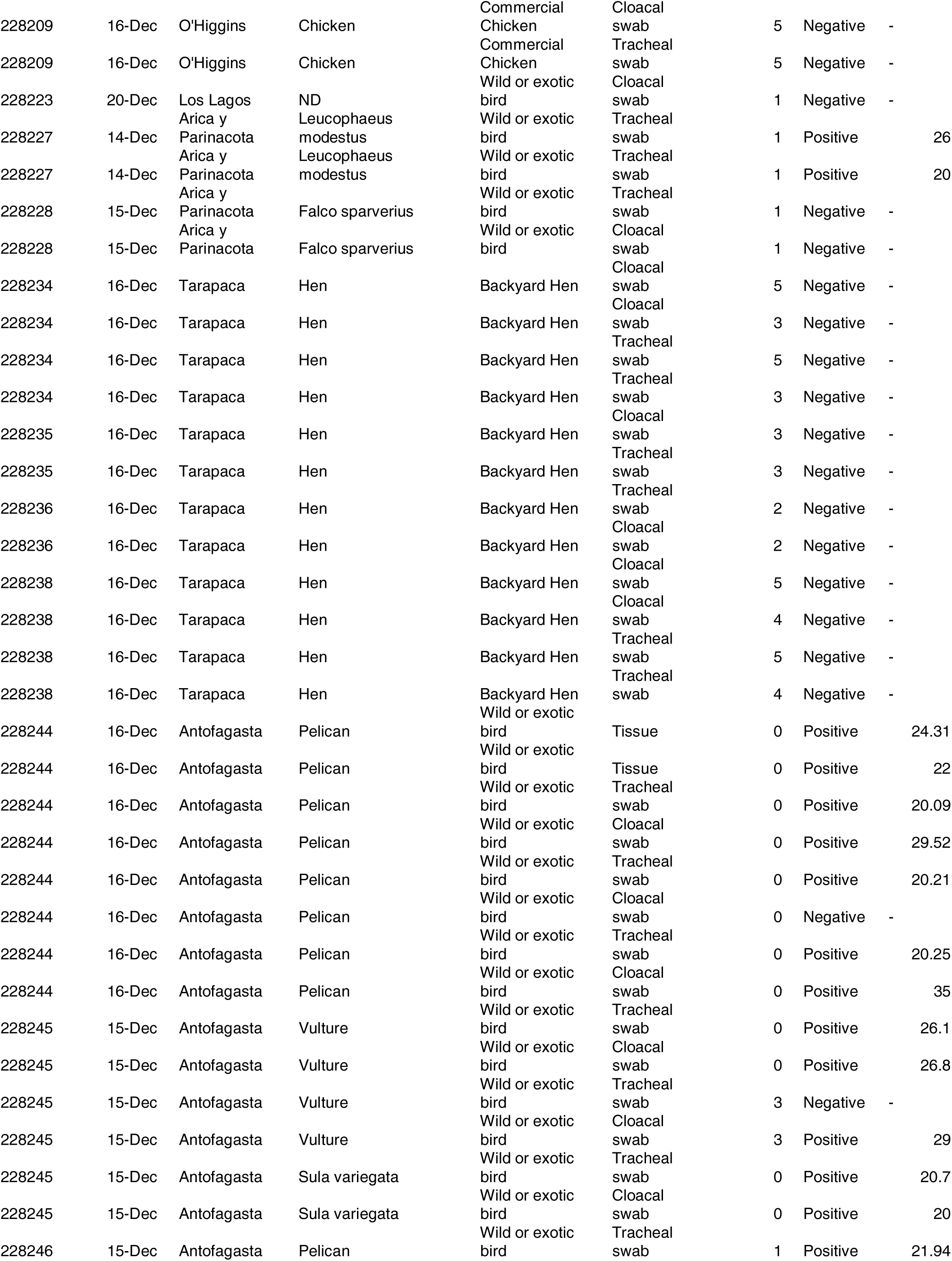
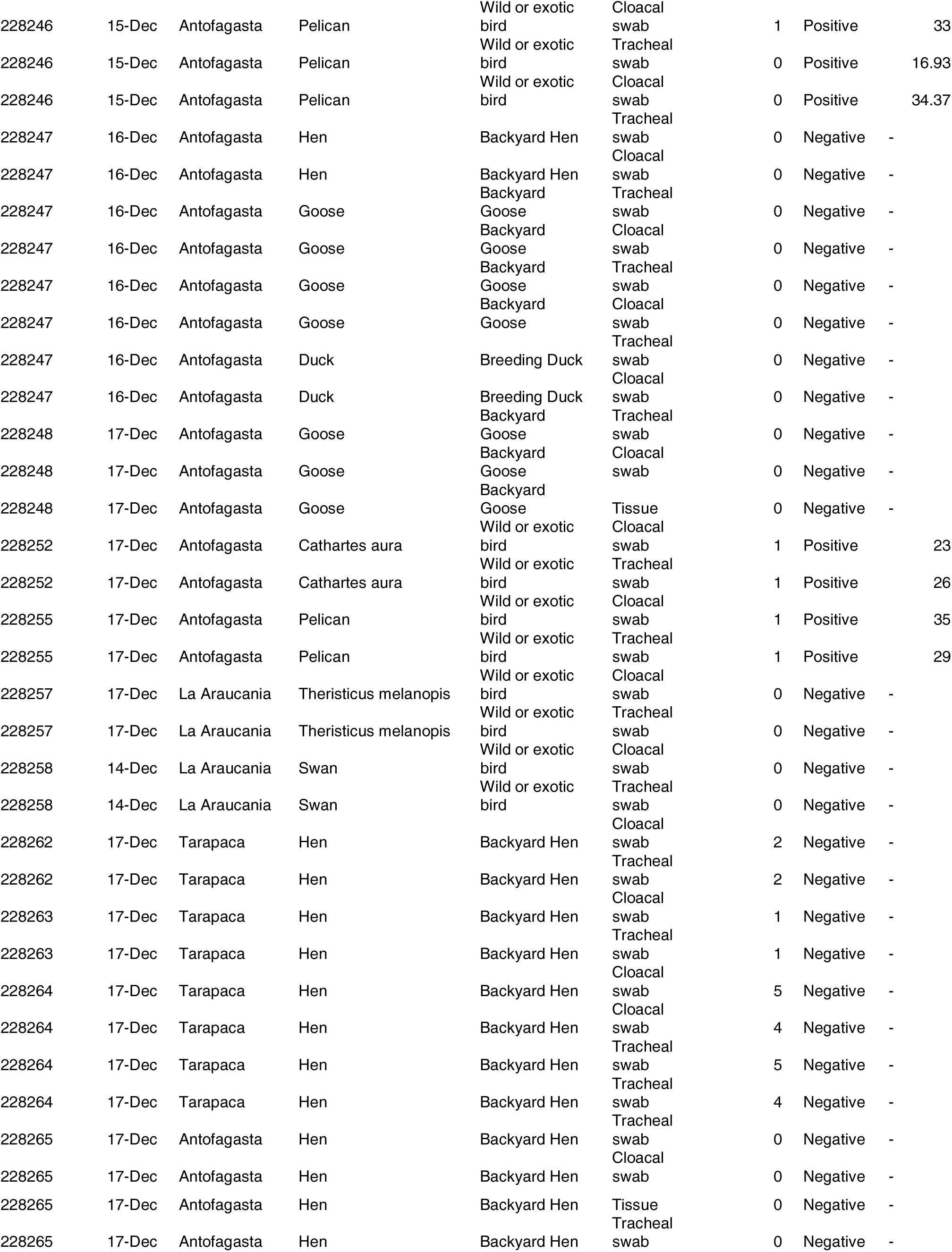
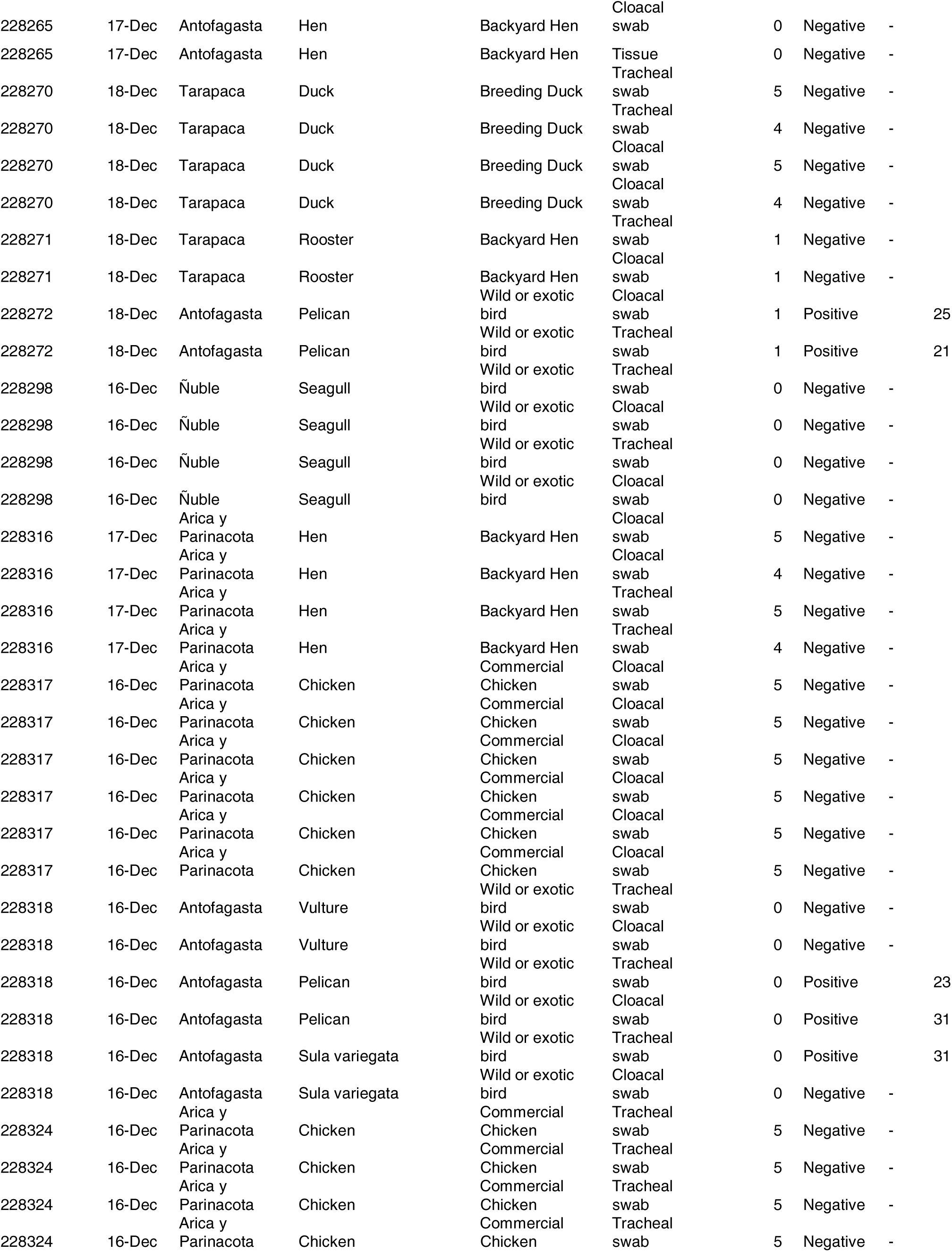
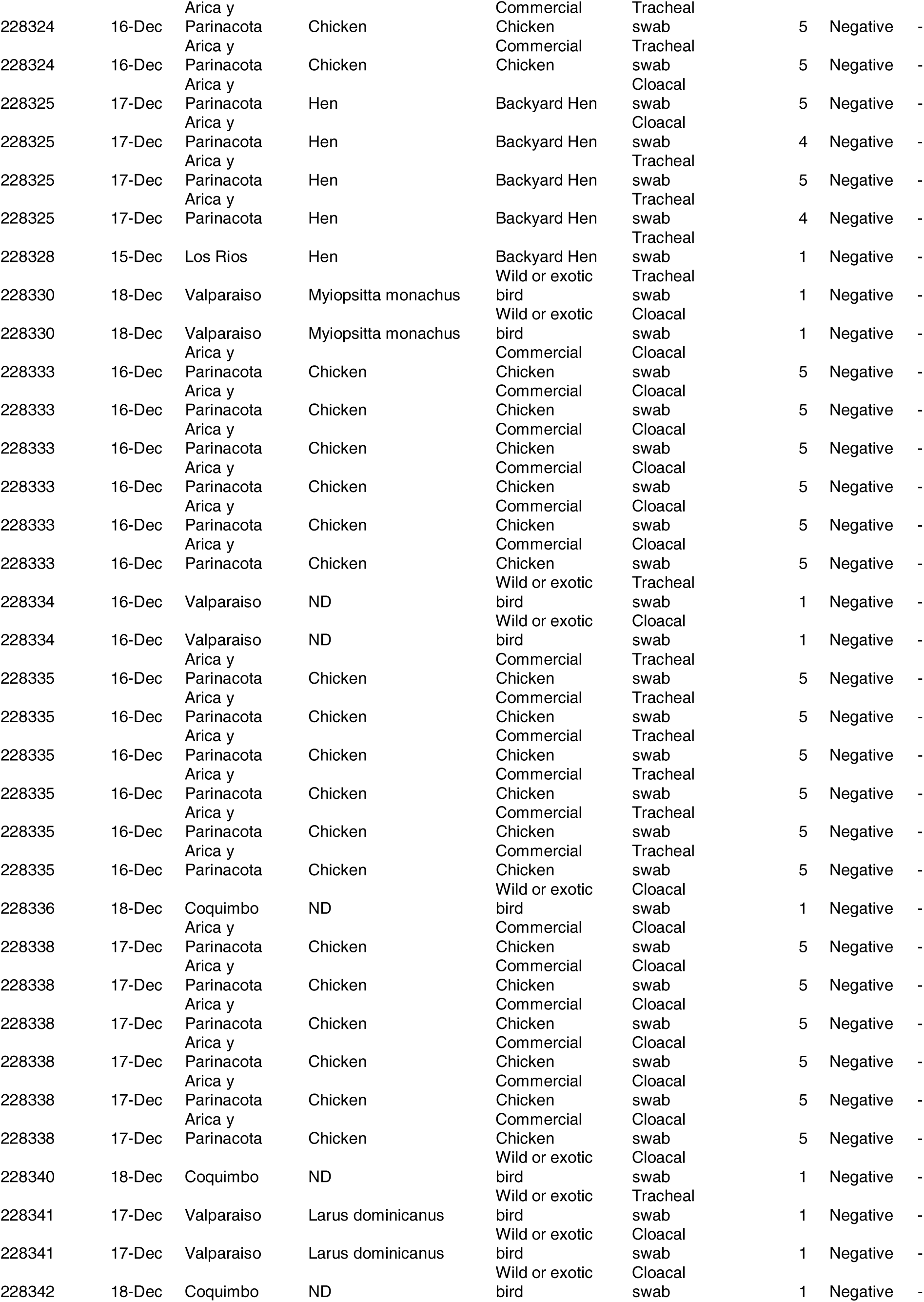
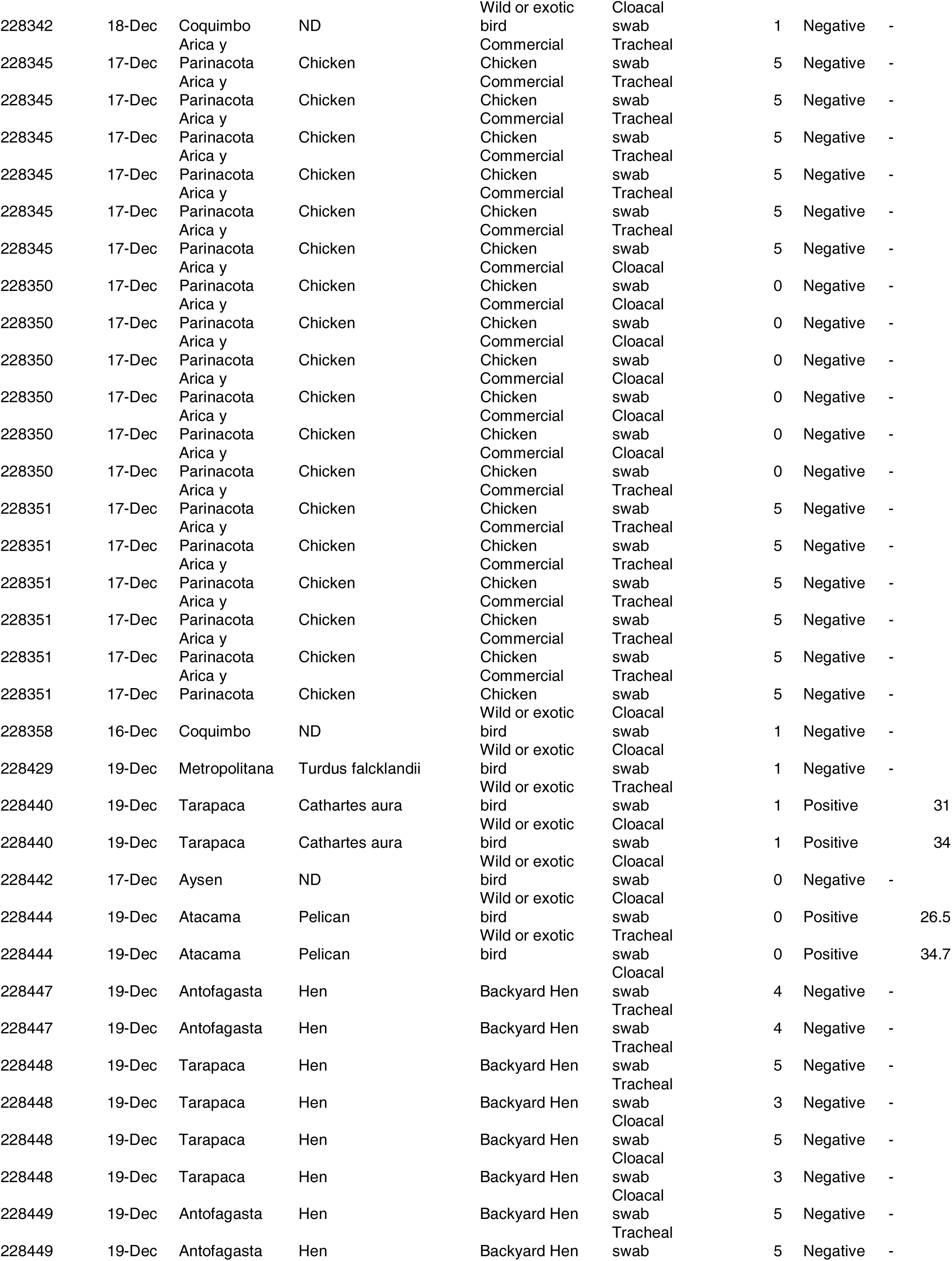
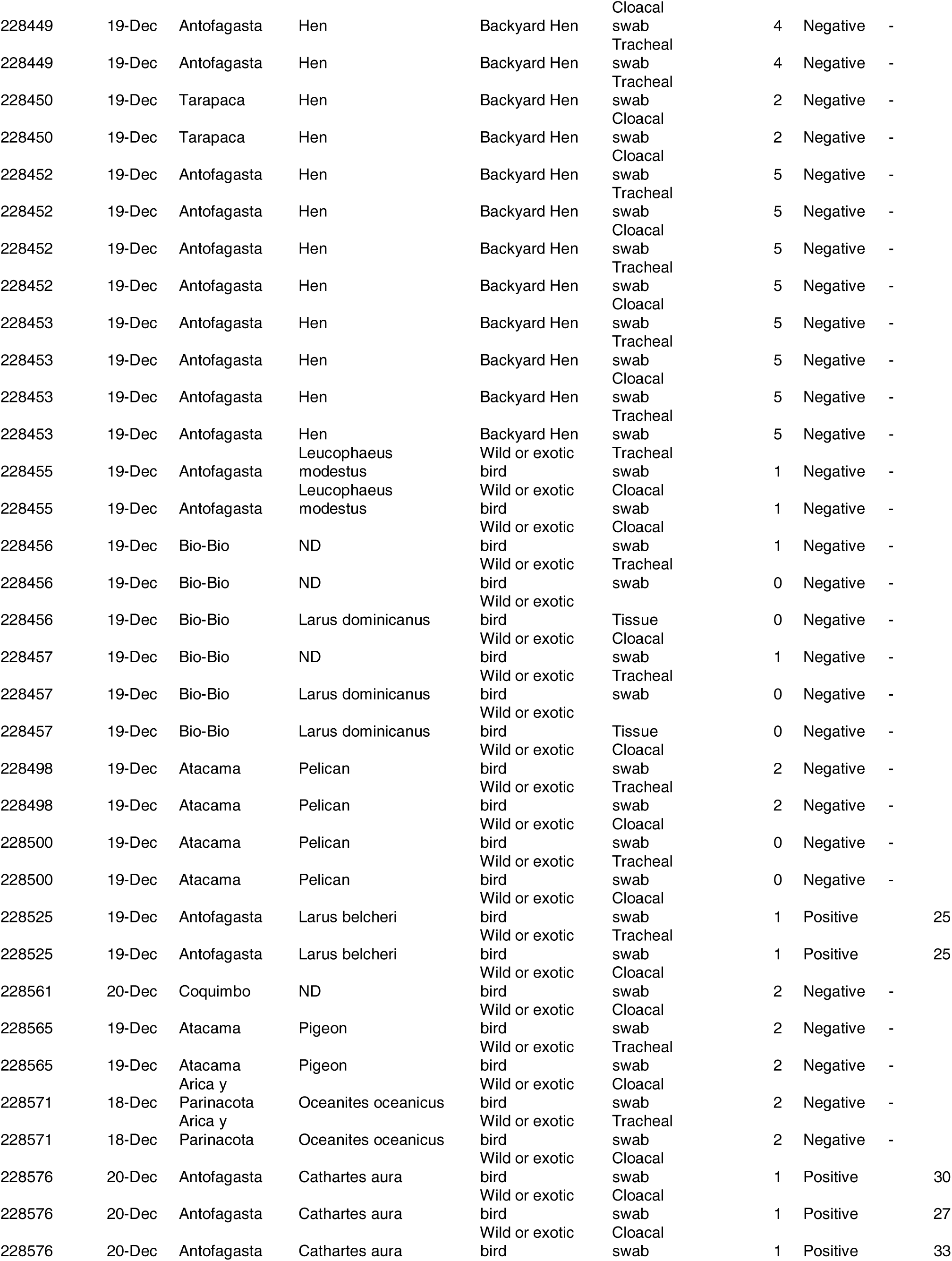
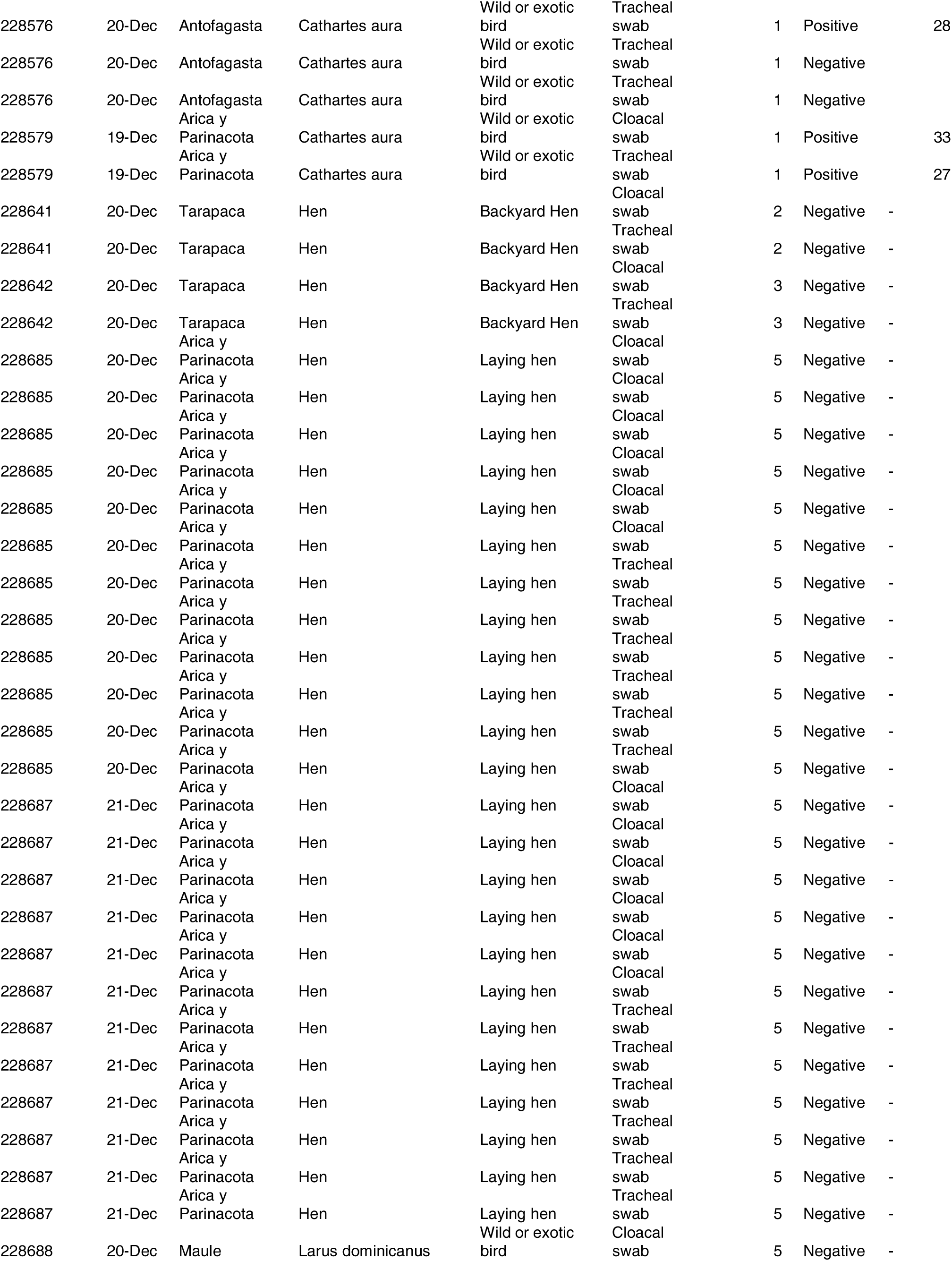
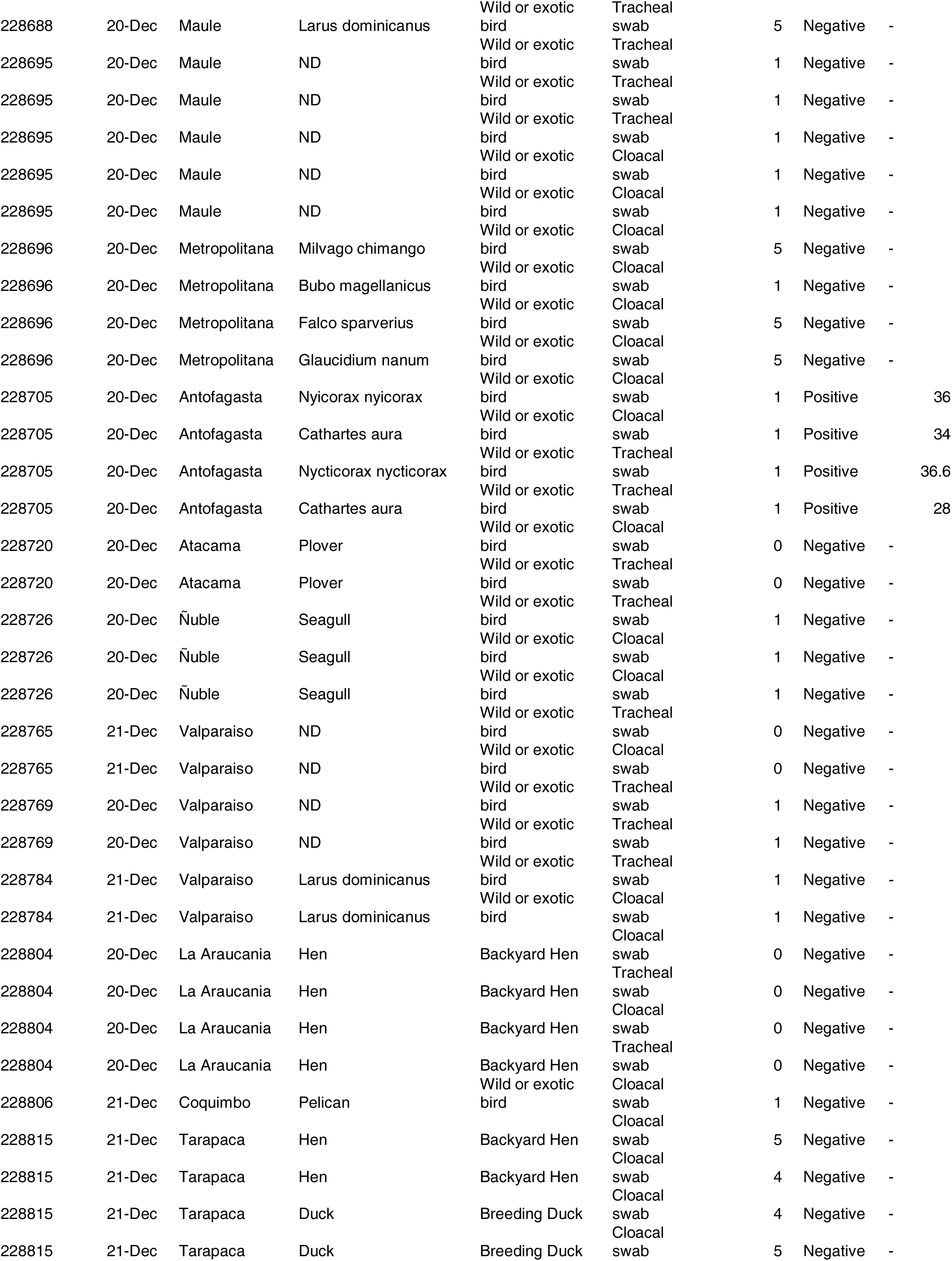
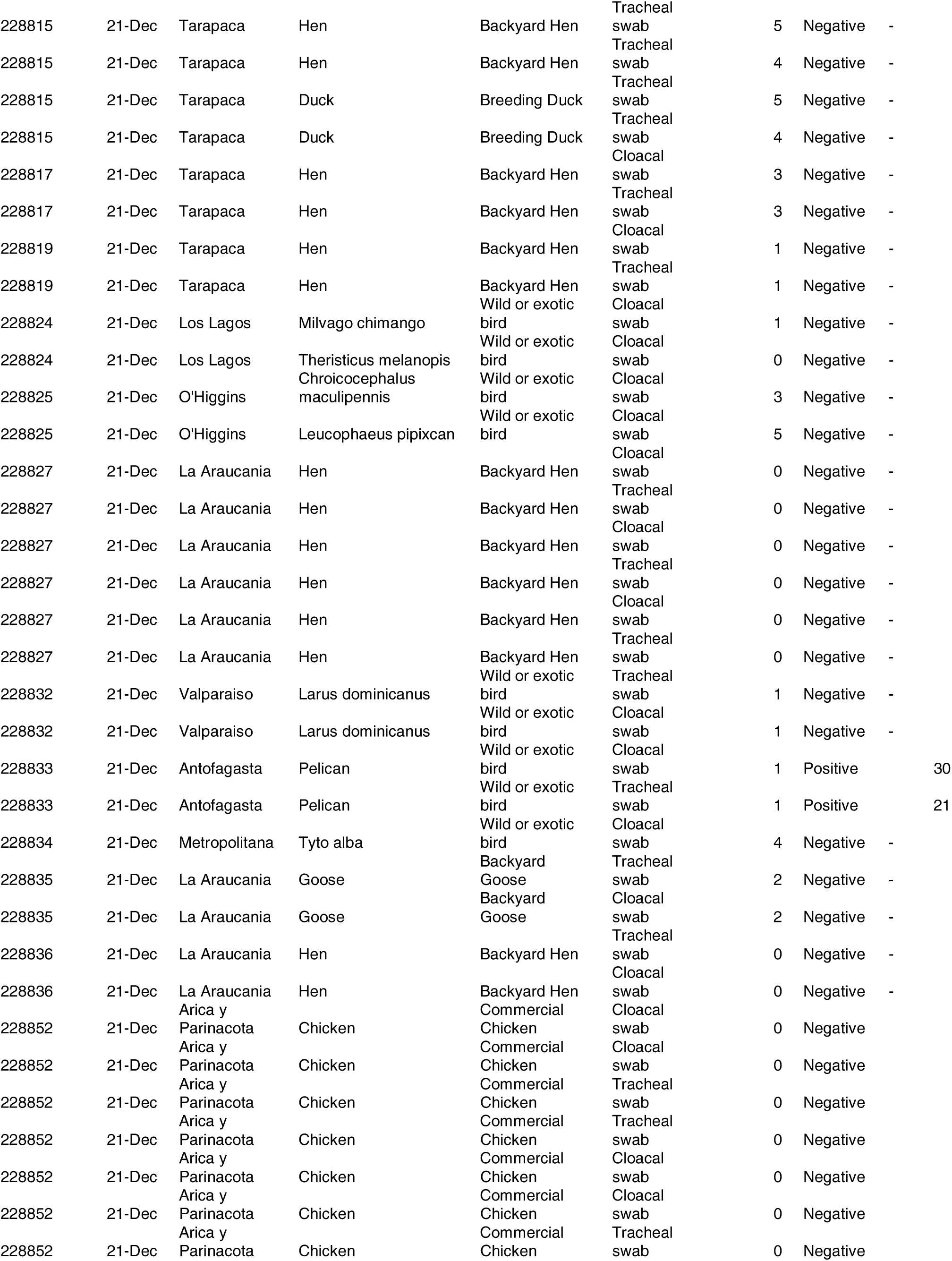
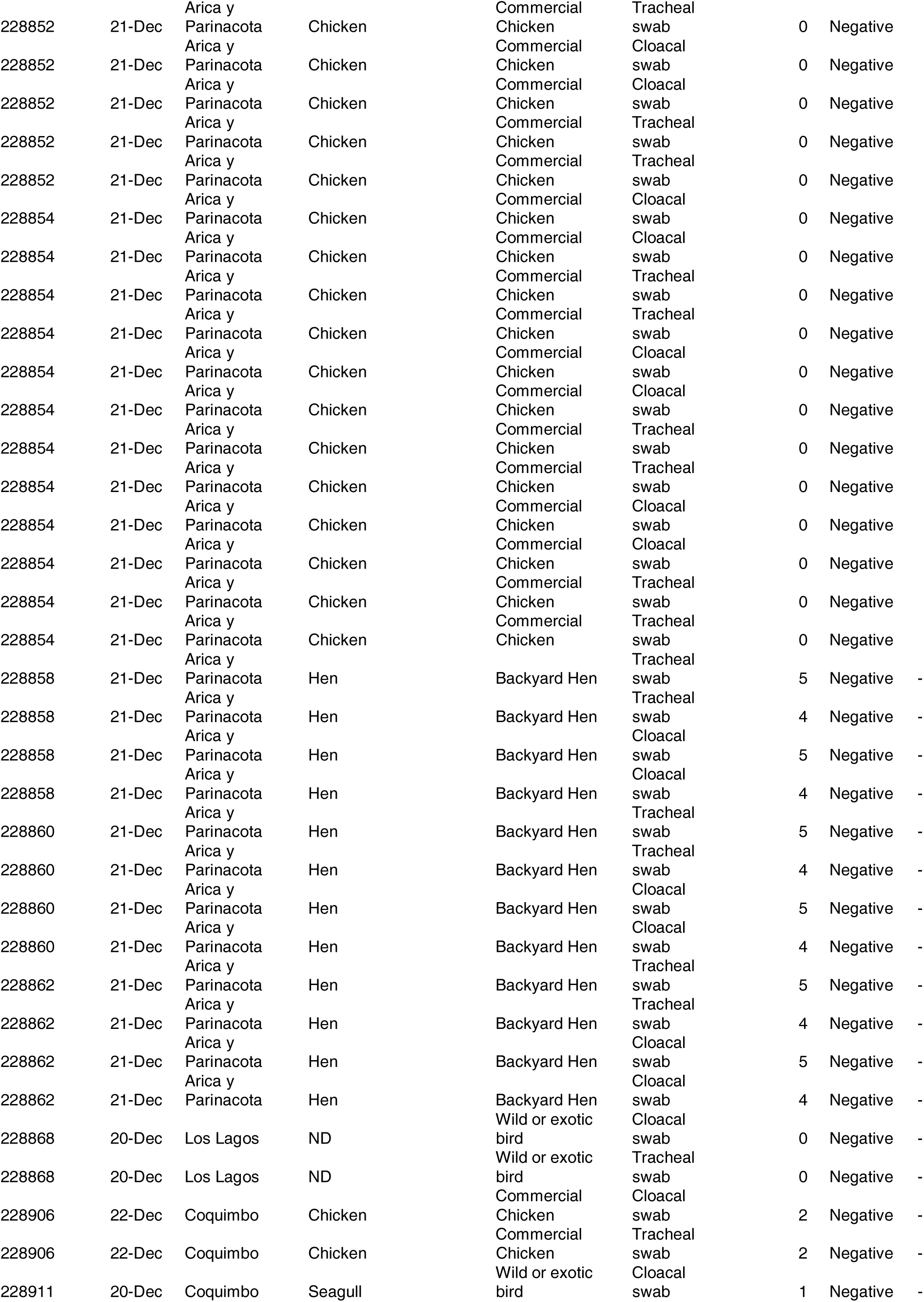
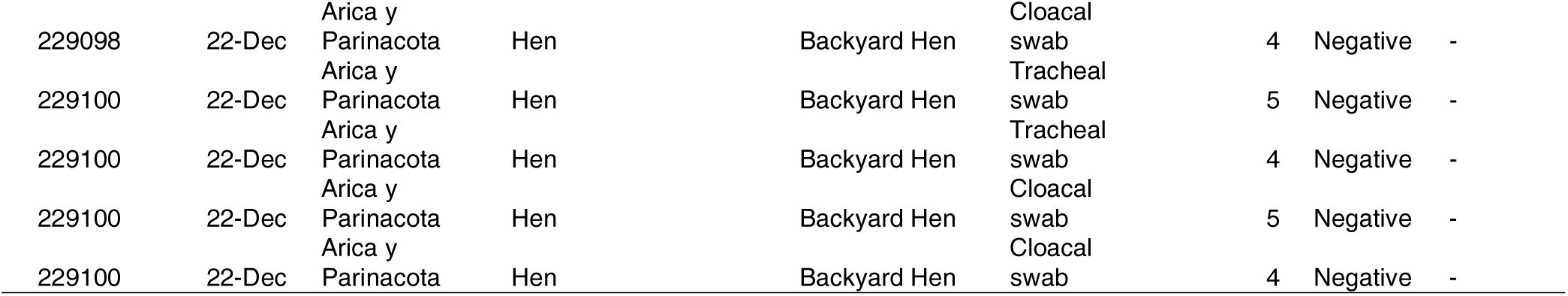
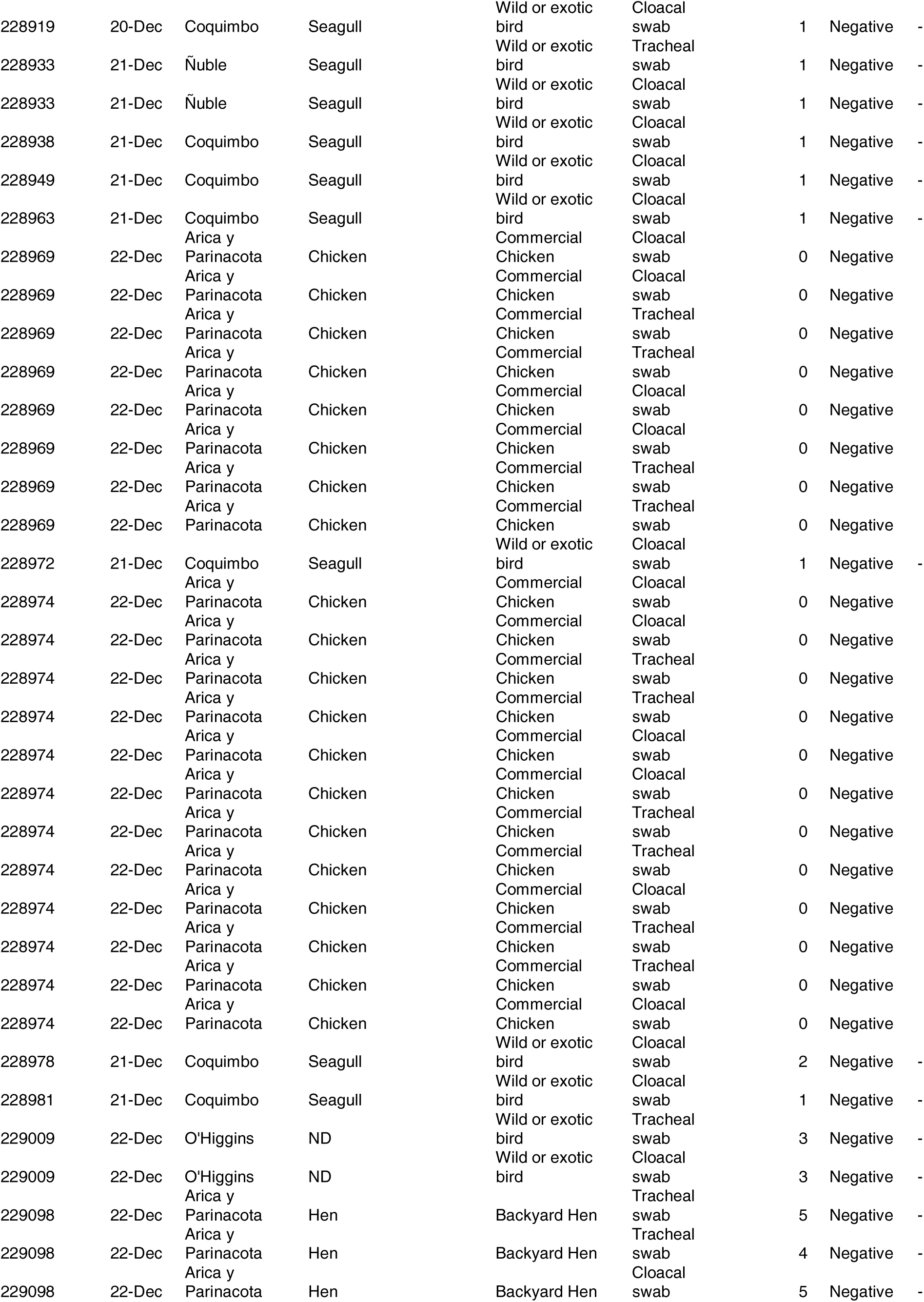
Samples of wild and domestic birds officially analyzed for HPAI detection by real time RT-PCR up to December 22, 2022, Chile.

**Appendix Table 2.**
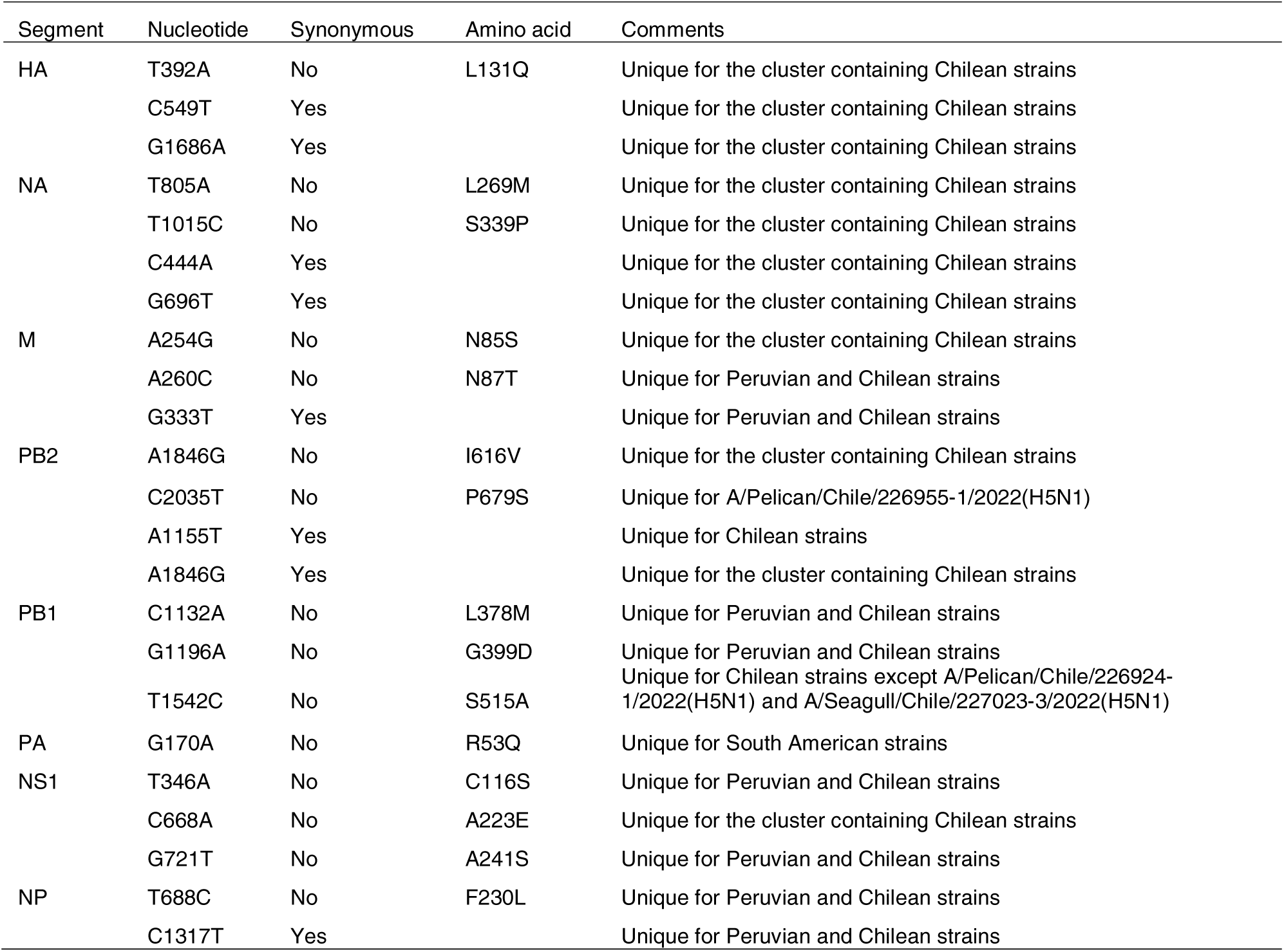
Featured mutations found in HPAI H5N1 Chilean viral strains sequenced in this study compared to representative strains included in the phylogenetic analyses from other countries.

**Appendix Figure 1.** Distribution of samples tested for avian influenza virus and affected pelicans are shown, Chile, December 22, 2022.

**Appendix Figure 2.** Maximum likelihood tree depicting the phylogeny of PB1 from isolates of H5N1 subtype 2.3.4.4b H5 clade. The Chilean-Peruvian subcluster is highlighted in red, the North American in blue, and other reference sequences in black.

**Appendix Figure 3.** Maximum likelihood tree depicting the phylogeny of PB2 from isolates of H5N1 subtype 2.3.4.4b H5 clade. The Chilean-Peruvian subcluster is highlighted in red, the North American in blue, and other reference sequences in black.

**Appendix Figure 4.** Maximum likelihood tree depicting the phylogeny of PA from isolates of H5N1 subtype 2.3.4.4b H5 clade. The Chilean-Peruvian subcluster is highlighted in red, the Venezuelan strains in green, Ecuadorian in pink, the North American in blue, and other reference sequences in black.

**Appendix Figure 5.** Maximum likelihood tree depicting the phylogeny of NP from isolates of H5N1 subtype 2.3.4.4b H5 clade. The Chilean-Peruvian subcluster is highlighted in red, the North American in blue, and other reference sequences in black.

**Appendix Figure 6.** Maximum likelihood tree depicting the phylogeny of M from isolates of H5N1 subtype 2.3.4.4b H5 clade. The Chilean-Peruvian subcluster is highlighted in red, the Venezuelan strains in green, the North American in blue, and other reference sequences in black.

**Appendix Figure 7.** Maximum likelihood tree depicting the phylogeny of NS from isolates of H5N1 subtype 2.3.4.4b H5 clade. The Chilean-Peruvian subcluster is highlighted in red, the North American in blue, and other reference sequences in black.

