## Supplementary figures and images for "Emergence and rapid dissemination of highly pathogenic avian influenza virus H5N1 clade 2.3.4.4b in wild birds, Chile"

### Appendix Figure 1

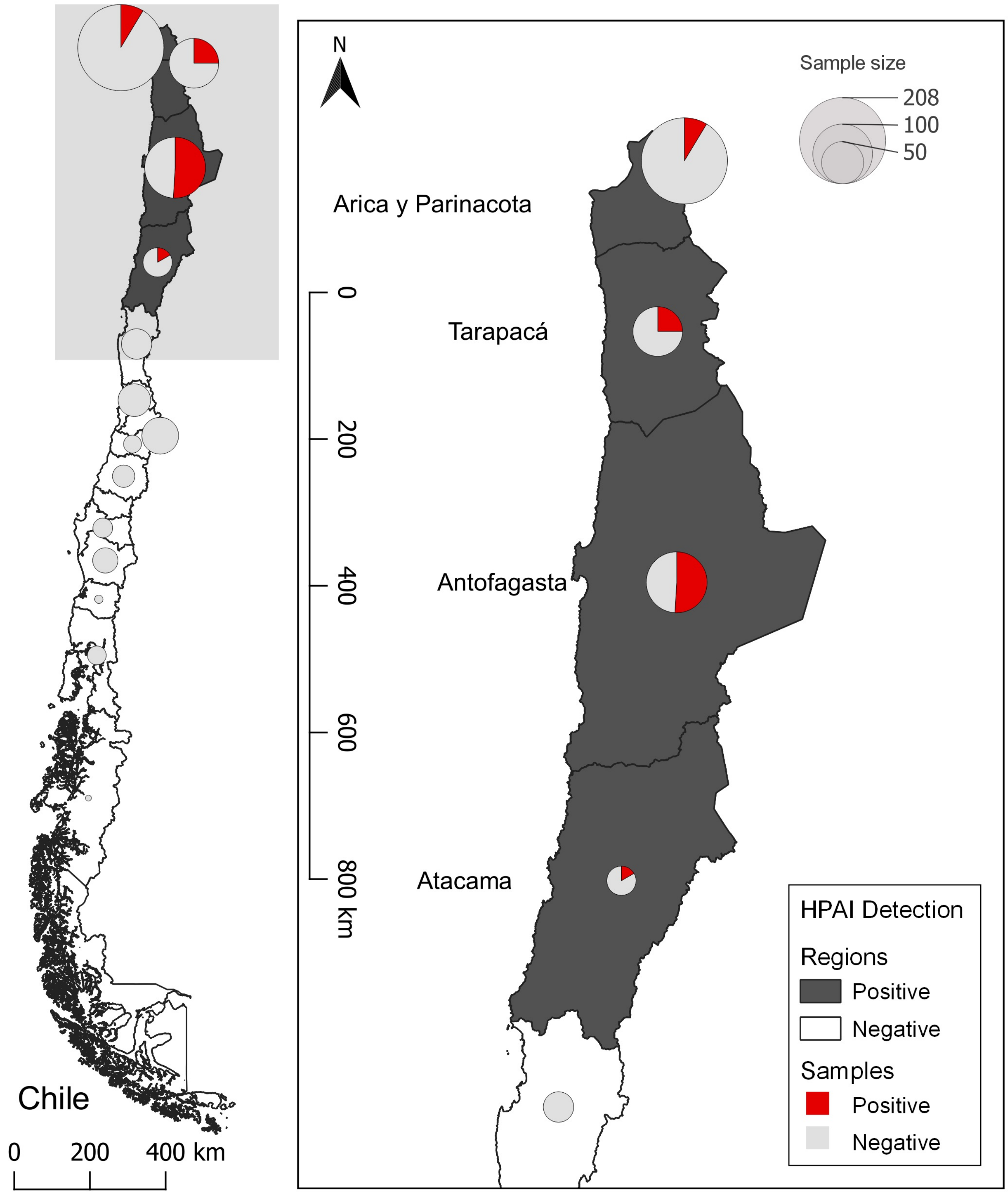

### Appendix Figures 2-7

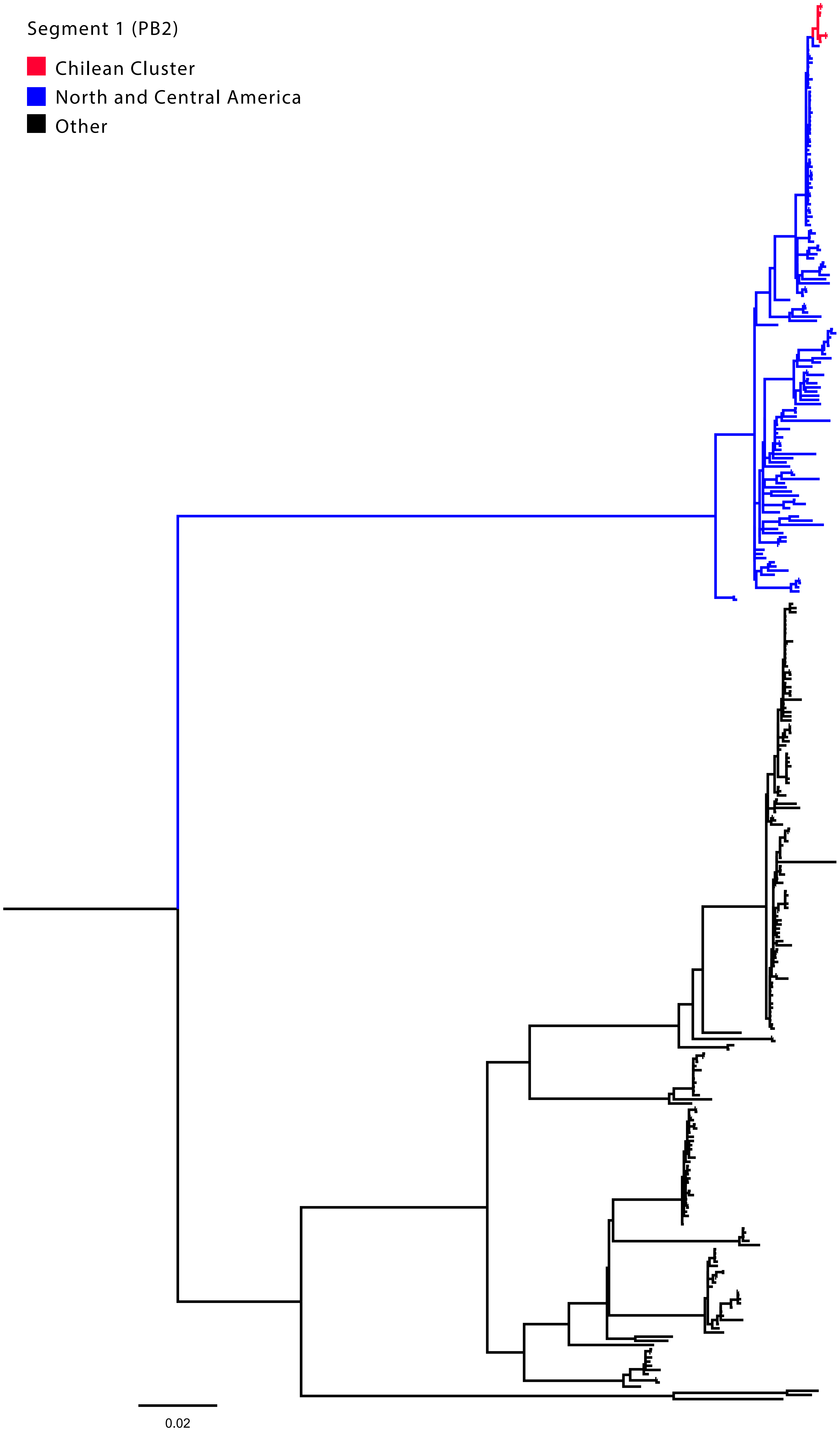

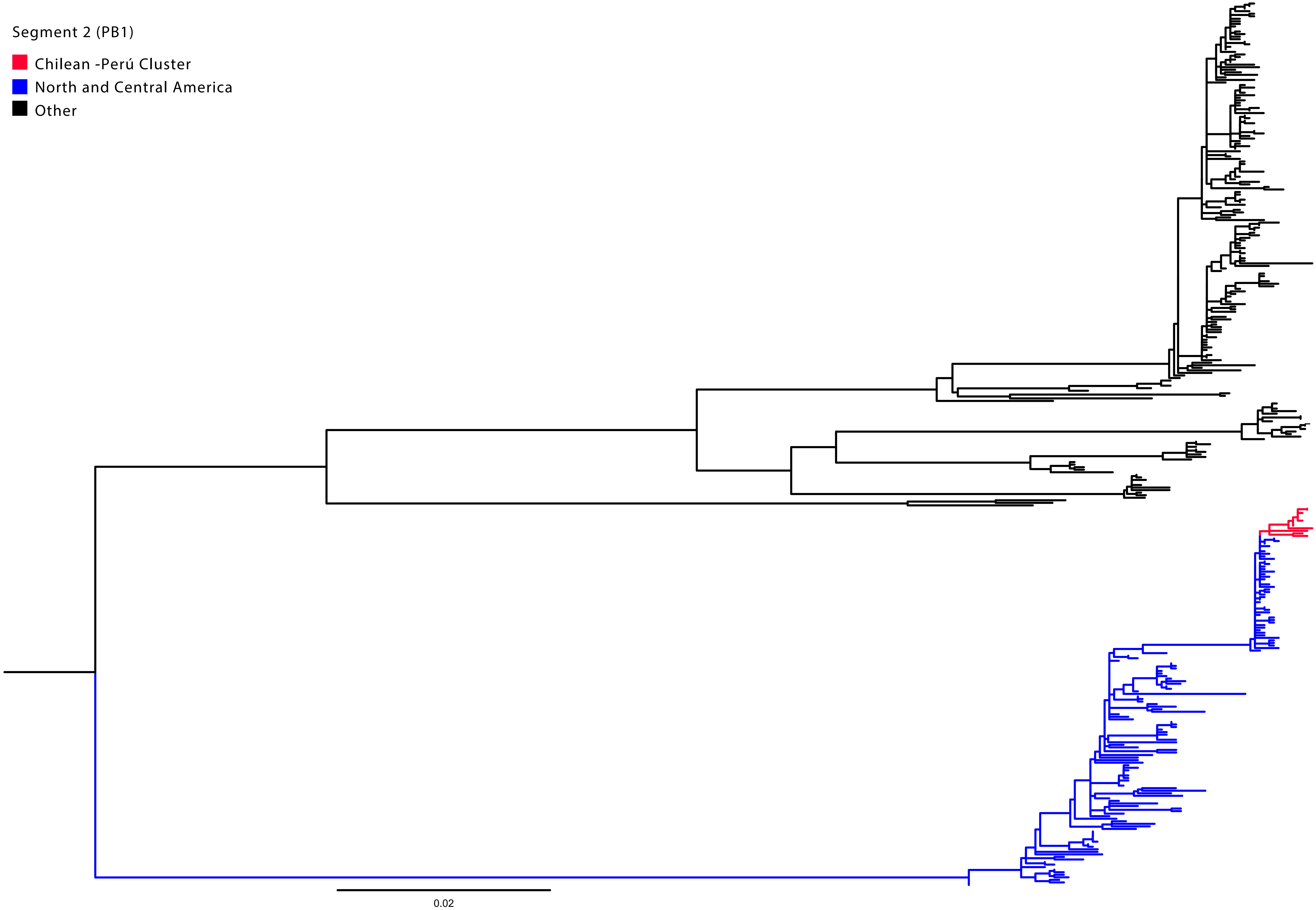

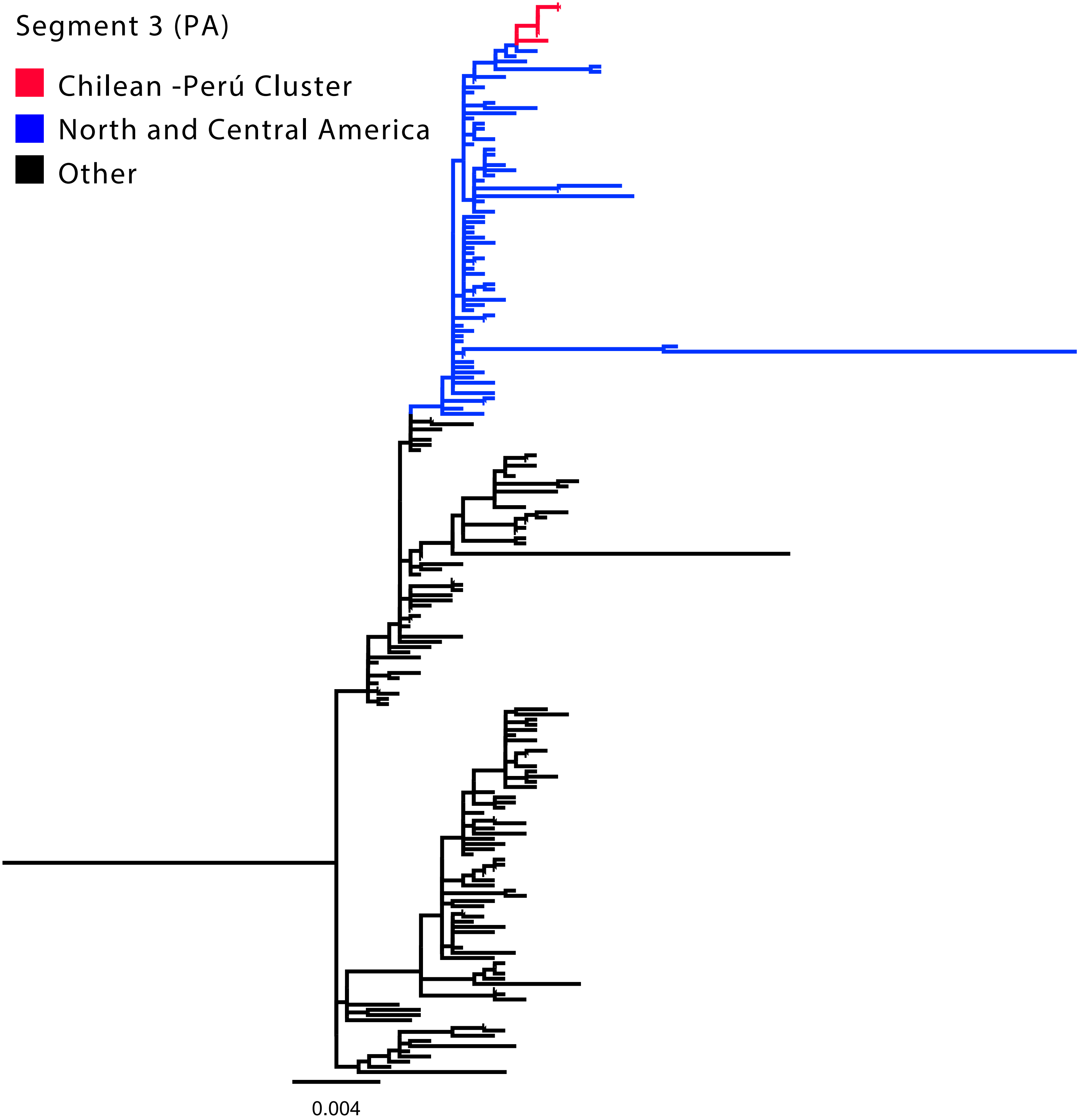

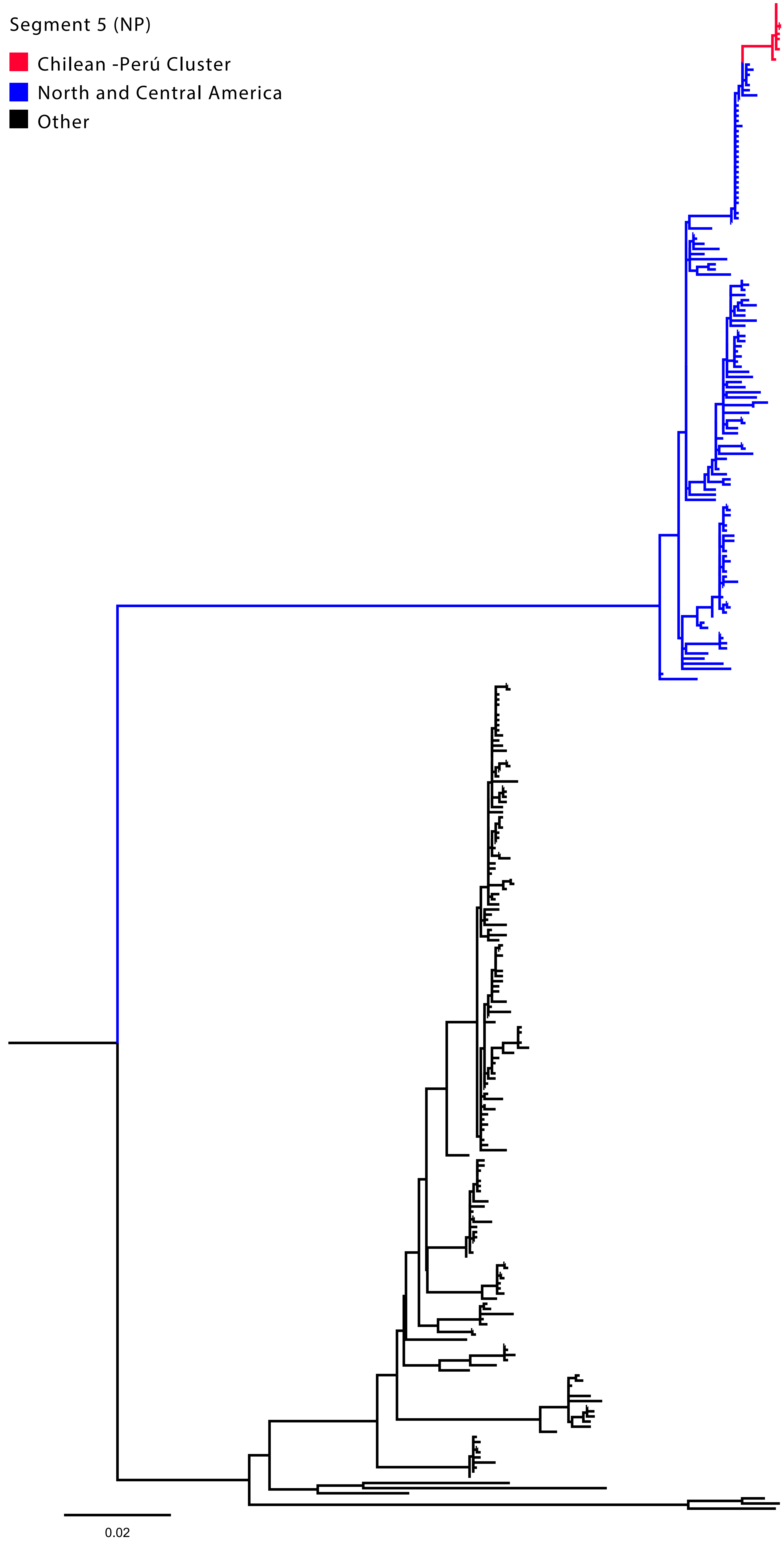

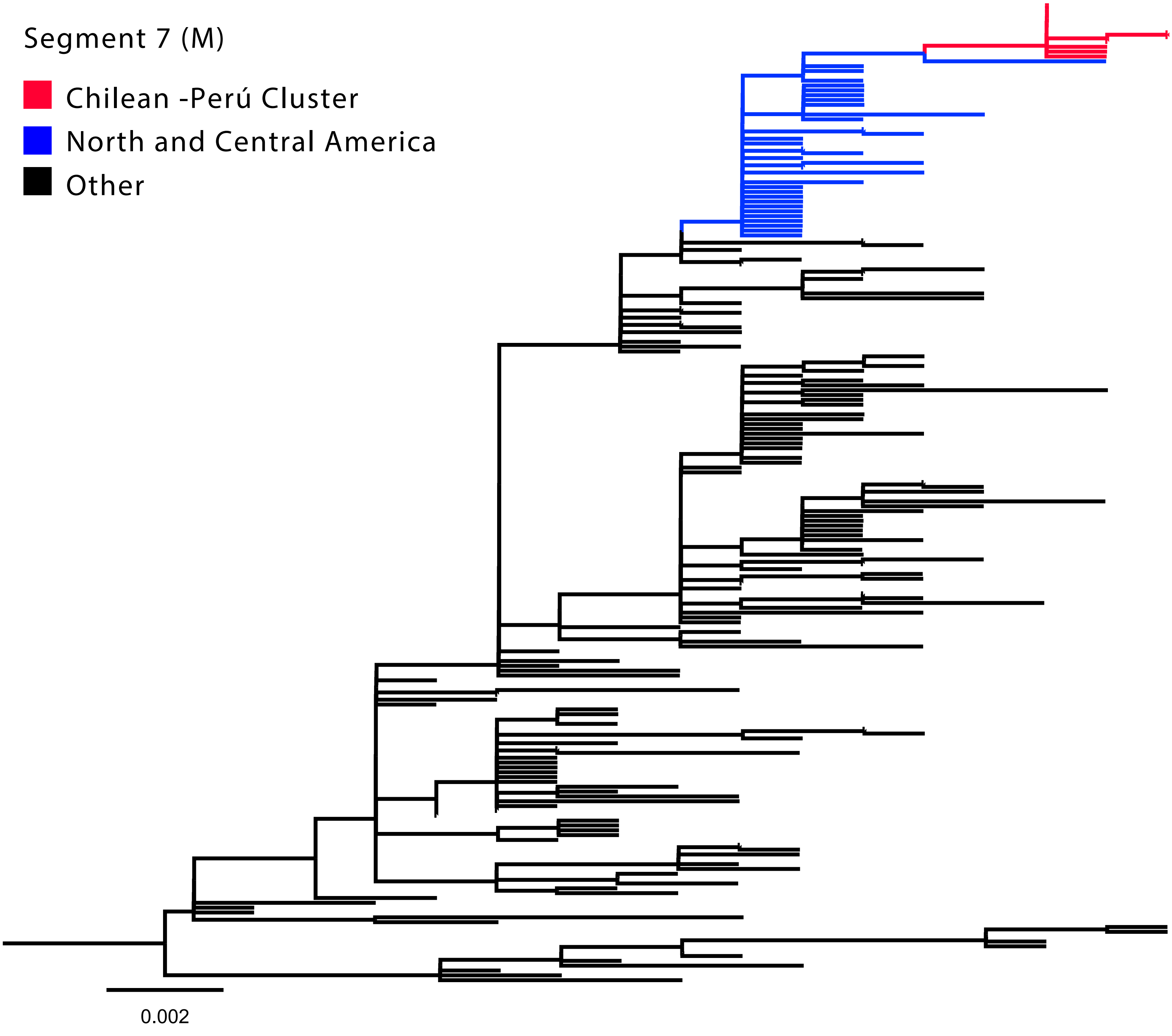

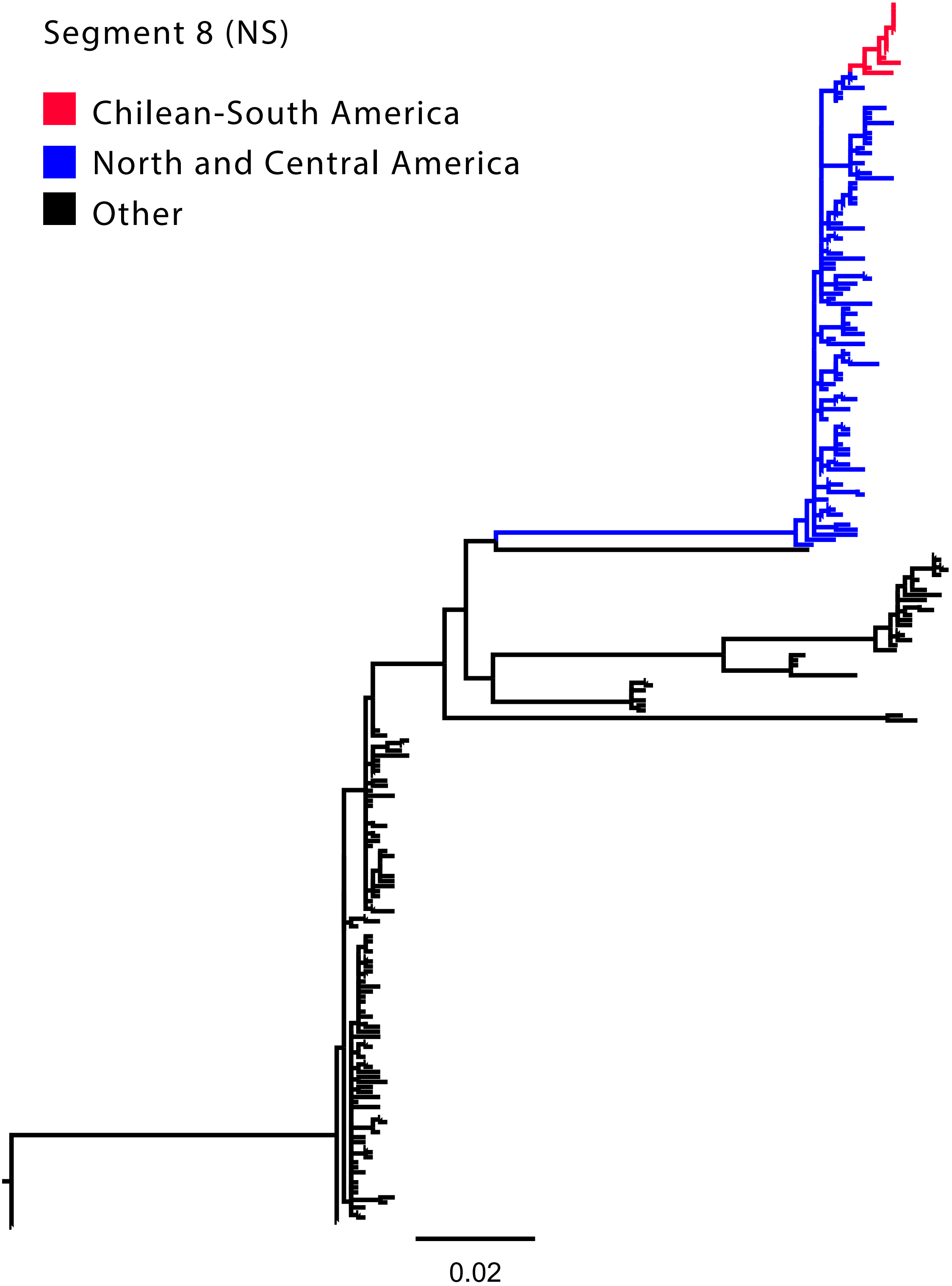
