## Appendix for "Emergence and rapid dissemination of highly pathogenic avian influenza virus H5N1 clade 2.3.4.4b in wild birds, Chile"

**Phylogenetic Analyses**

Datasets for each influenza viral segment sequences were created by retrieving reference sequences from NCBI(1) and GISAID databases (2). The sequences were aligned using MUSCLE (3) in Mega X (4). Duplicate sequences were removed, and the phylogeny was estimated using the maximum likelihood method available in IQ-TREE v2.1.2, employing -m TEST option for nucleotide substitution model selections and 1000 ultrafast bootstrap replications in CIPRES (5,6). Each phylogenetic tree was visualized using Figtree (http://tree.bio.ed.ac.uk/software/figtree/). Also, we performed Bayesian evolutionary analysis by sampling trees (BEAST) analysis for the HA and NA segment(7).
